# Construction, Deployment, and Usage of the Human Reference Atlas Knowledge Graph for Linked Open Data

**DOI:** 10.1101/2024.12.22.630006

**Authors:** Andreas Bueckle, Bruce W. Herr, Josef Hardi, Ellen M. Quardokus, Mark A. Musen, Katy Börner

## Abstract

The Human Reference Atlas (HRA) for the healthy, adult body is being developed by a team of international, interdisciplinary experts across 20+ consortia. It provides standard terminologies and data structures for describing specimens, biological structures, and spatial positions of experimental datasets and ontology-linked reference anatomical structures (AS), cell types (CT), and biomarkers (B). We introduce the HRA Knowledge Graph (KG) as central data resource for HRA v2.2, supporting cross-scale, biological queries to Resource Description Framework graphs using SPARQL. In February 2025, the HRA KG covered 71 organs with 5,800 AS, 2,268 CT, 2,531 B; it had 10,064,033 nodes, 171,250,177 edges, and a size of 125.84 GB. The HRA KG comprises 13 types of Digital Objects (DOs) using the Common Coordinate Framework Ontology to standardize core concepts and relationships across DOs. We (1) provide data and code for HRA KG construction; (2) detail HRA KG deployment by Linked Open Data principles; and (3) illustrate HRA KG usage via application programming interfaces, user interfaces, data products. A companion website is at https://cns-iu.github.io/hra-kg-supporting-information.

## Background & Summary

The multimodal, three-dimensional (3D) Human Reference Atlas (HRA)^1^ aims to map the healthy, adult human body across scales—from the whole body to the single cell and biomarker levels. Data from different sources (organs, technologies, and labs), many built with standard operating procedures (SOPs, https://humanatlas.io/standard-operating-procedures), need to be integrated. The HRA Knowledge Graph (KG) defines and provides the core data structures that are used to store, link, and query HRA data.

KGs are widely used to interlink data about relevant entities within a specific domain or task. The Google Knowledge Graph (https://blog.google/products/search/introducing-knowledge-graph-things-not) supports Google Search with its billions of searches processed daily. Major online shopping retailers such as Amazon^2^ use knowledge graphs to organize products, searches, and media items^3^. KGs across domains are structured using vocabularies, e.g., Friend of a Friend (FOAF, http://xmlns.com/foaf/spec), Simple Knowledge Organization System (SKOS, https://www.w3.org/2004/02/skos), and Music Ontology (http://musicontology.com). Plus, there exist collaborative efforts for publishing structured data on the web. For example, https://schema.org promotes the structured representation for data on the web and is used in applications from Google, Microsoft, and Pinterest to create data-driven web experiences. An overview of other commonly used vocabularies is available on https://www.w3.org/wiki/TaskForces/CommunityProjects/LinkingOpenData/CommonVocabularies.

### Summary

In this paper, we present the HRA KG v2.2, which uses 10 ontologies to interlink 33 anatomical structures (AS), cell types (CT), plus biomarkers (B) tables (see **Box 1**), 71 3D Reference Objects for organs, 22 Functional Tissue Units (FTUs)^4^, 11,698 single-cell (sc) datasets, and other HRA Digital Objects (DOs). The HRA KG is accessible via (1) SPARQL, (2) the RESTful HRA API with clients available in JavaScript, TypeScript, Angular 17+, and Python 3.6+ ((https://humanatlas.io/api), and (3) several interactive user interfaces (UIs).. Specifically, we present open code and infrastructure to construct the HRA KG out of disparate data across tabular/non-tabular and nested/flat HRA DOs while ensuring processed data conforms to 5-star Linked Open Data (LOD)^5^ principles. The HRA KG data can be accessed via content negotiation from https://lod.humanatlas.io and dynamically queried via its SPARQL (see **Box 1**) endpoint at https://lod.humanatlas.io/sparql. This can be used to obtain any version of any DO in any supported Resource Description Framework (RDF, see **Box 1**) format hosted by the HRA KG.

##### Key technologies used for constructing, deploying, and using the HRA KG

- **3D Reference Objects:** Mesh-based 3D models describing organs in the male or female body of the HRA. They are used in HRA applications to register and explore tissue blocks and associated datasets^6^. All 71 3D Reference Objects of HRA v2.2 have crosswalks that link individual 3D AS to ontology terms in Uberon^7^ or Foundational Model of Anatomy (FMA)^8,9^.
- **Anatomical structures (AS), cell types (CT), plus biomarkers (B) tables (ASCT+B):** ASCT+B tables are authored by multiple experts across many consortia. They capture the relationship between AS (and the AS located in them), CT found inside these AS, and the B (genes, proteins) used to characterize the CT, see details in related publications^1,10^.
- **Cell Type Annotation (CTann):** Azimuth^11^, CellTypist^12,13^, and popV^14^ are used to assign CT to cells from sc/snRNA-seq studies. Manually compiled crosswalks are used to assign ontology IDs to CTann CT, see details in a related publication^1^.
- **Common Coordinate Framework (CCF) Ontology**: The CCF Ontology^15^ provides the main vocabulary for constructing atlases of the human body, including the HRA. Critically, the CCF provides the framework for constructing atlases, but is not an atlas itself. It includes concepts and properties needed to describe the human body, from organs down to CT and B, for organizing spatial data, and for capturing donor, sample and dataset metadata published in the KG.
- **Content negotiation**: This mechanism is used by HTTP servers to serve different versions of a resource at the same Uniform Resource Identifier (URI) based on the parameters given in the HTTP request (https://developer.mozilla.org/en-US/docs/Web/HTTP/Content_negotiation). HTTP requests can specify headers which provide additional information for the server to act on. Using the *Accept* header, an agent can specify what format (or a ranked list of acceptable formats) they would like the response returned in. A web browser will typically request *text/html*, but machines or programmers may request other formats like *application/json* or any of the RDF formats supported by the HRA KG.
- **Crosswalks:** An ontological matching of terms in HRA DOs to ontology terms in the ASCT+B tables^10^. Crosswalks can link, e.g., 2/3D Reference Objects of organs to AS and CT, and OMAPs^16^ to CT and B. This definition is adapted from a related publication^1^.
- **HRA Digital Objects (DOs):** HRA DOs are the data components for generating the HRA KG. They are explained in detail in the Methods section. ASCT+B tables, 3D Reference Objects, and OMAPs are examples of HRA DOs that are processed to become part of the HRA KG.
- **Linked Open Data (LOD)**: A common data sharing pattern^17^, developed for the Semantic Web (https://www.w3.org/2001/sw/wiki/Main_Page) that describes how to structure and share semantically rich data that allows for maximum reuse and utility. To be LOD, the data should have an open license, use URIs in the data to name entities whose URIs resolve (i.e., can be queried either directly via web request or via SPARQL) to retrieve structured data in RDF about that URI and link to other resources via URIs.
- **Linked Data Modeling Language (LinkML)**: LinkML (https://linkml.io/linkml/)^18^ is a flexible linked data modeling language that allows us to author schemas in YAML (https://yaml.org) which describe the structure of one’s data. Additionally, it is a framework for working with and validating data in a variety of formats (JavaScript Object Notation [JSON], RDF, tab-separated values [TSV]), with generators for compiling LinkML schemas to other frameworks.
- **Organ Mapping Antibody Panels (OMAPs)**: Tabular data structures with panels of experiment-derived and tested antibodies to target proteins for identifying AS, CT, cell states, or cell membrane staining in organs.
- **Persistent Uniform Resource Locator (PURL)**: A PURL is a type of URL pointing to a resolution service rather than to a website. This enables the resolution service to use content negotiation to determine what content is needed (e.g., HTML for humans, structured data for machines) and to redirect or directly return the relevant data. PURLs are used in LOD to provide persistent, resolvable URIs for entities so that they can be referenced without worry of the URIs changing.
- **Resource Description Framework (RDF)**: RDF is a standard to represent connected data on the web. It defines relationships between data objects, enabling exchange of structured information through triples consisting of a subject, predicate, and object (https://www.w3.org/RDF/).
- **SPARQL Protocol and RDF Query Language**: A query language for RDF graphs (https://www.w3.org/TR/sparql11-query), SPARQL can be used to write declarative code to retrieve triples that describe two entities and their relationship in an RDF graph. Triples have a subject-predicate-object relationship.
- **Subject matter experts (SMEs)**: Individuals who possess specialized training in areas related to HRA construction, such as anatomists, surgeons, clinicians, and physicians. SMEs may have valuable knowledge about entire organ systems, individual organs, or parts thereof, such as their cellular or molecular make-up, or have expertise in experimental procedures.

### Ontologies and KGs

Linking HRA DOs (see **Box 1**) to ontologies and, by extension, wider domains of expert knowledge is a high priority for HRA KG construction. In the biomedical domain, ontologies are widely used to structure data, which is of high relevance to HRA KG construction. For example, the National Center for Biomedical Ontology (NCBO) BioPortal provides easy access to 1,168 ontologies and the EMBL-EBI Ontology Lookup Service (OLS) supports 267 ontologies. The Uber-anatomy ontology (Uberon, https://www.ebi.ac.uk/ols4/ontologies/uberon)^7^ is a cross-species ontology representing body parts, organs, and tissues, primarily focused in vertebrates. The Cell Ontology (CL, https://obofoundry.org/ontology/cl.html)^19^ is also a cross-species ontology, but it focuses on classifying and describing cells. These two ontologies are linked data, so we can determine assertions such as *kidney cortical cell* (http://purl.obolibrary.org/obo/CL_0002681) is *part of* (http://purl.obolibrary.org/obo/BFO_0000050) *cortex of kidney* (http://purl.obolibrary.org/obo/UBERON_0001225) from the knowledge represented in the ontologies. Ontologies are an indispensable part of generating, using, and maintaining KGs as they enable unifying nomenclature across assay types, organs, donors, teams, and consortia. A recent publication by He et al.^20^, featuring the HRA, shows how ontologies can be used to model, integrate, and reason over previously siloed clinical, pathological, and molecular kidney data for precision medicine. It highlights the development of the precision medicine metadata ontology (PMMO) to integrate dozens of variables between the Kidney Precision Medicine Project (KPMP, https://www.kpmp.org)^21,22^ and Chan Zuckerberg Initiative (CZI) CELLxGENE (CxG) data (https://cellxgene.cziscience.com). It then shows specific use cases in detecting healthy vs. acute kidney infection (AKI)/chronic kidney disease (CKD) disease states in cells supported by PMMO, Kidney Tissue Atlas Ontology (KTAO), and the HRA’s CCF Ontology, described in a related publication^15^.

Biomedical KGs use and interlink multiple ontologies to store and query data. For example, the Unified Medical Language System (UMLS)^23^ “metathesaurus” contains approximately 3.4 million biomedical concepts, updated every 6 months in May and November and is derived from other biomedical terminologies and ontologies. The Petagraph KG^24^ uses the UMLS metathesaurus to integrate biomolecular datasets and connects them to approximately 200 cross-referenced ontologies to support exploration of gene variant epistasis as well as biological assertions with reduced dimensionality, and to link relevant features to chromosome position and chromosomal neighborhoods. The Human BioMolecular Atlas Program (HuBMAP, https://hubmapconsortium.org)^25,26^ Unified Biomedical Knowledge Graph^1^ (UBKG, https://ubkg.docs.xconsortia.org and https://github.com/x-atlas-consortia/ubkg-neo4j) connects HuBMAP experimental data to ontologies. The Scalable Precision Medicine Open Knowledge Engine (SPOKE, https://spoke.ucsf.edu)^27,28^ processes 41 databases (53 million edges) and 11 ontologies to create an integrated graph with user access via a Representational State Transfer (REST) application programming interface (API). Petagraph, HuBMAP, and SPOKE use the Neo4J graph platform (https://neo4j.com). Efforts like BioCypher (https://biocypher.org)^29^ enable the rapid construction and maintenance of KGs at lower cost. This also addresses the lack of reusability and integrability, where KGs are built manually for a specific task, and, as a result, do not adhere to Findable, Accessible, Interoperable and Reusable (FAIR)^30^ principles. KGs can be used to extract knowledge across constantly evolving ontologies and data in various states of accessibility (private and public).

### Updates since the CCF Ontology paper

The specimen, biological structure, and spatial ontologies in support of a HRA (v1.2) using CCF v2.0.1 were introduced in a prior publication^15^. Starting with HRA v2.0, published in December 2023, the CCF Ontology (v3.0) is separated from the HRA collection. CCF v3.0 is a DO of type (*vocab*) and the HRA collection is a DO of type (*collection*). Before then, it was a graph, essentially the HRA *collection* plus the CCF Ontology. Now the HRA collection references the CCF and is compiled from a collection of curated HRA DOs and is hosted by the HRA KG at (https://purl.humanatlas.io/collection/hra/v2.2). This change was necessary to establish a boundary between the framework for creating atlases, the CCF, and a specific atlas, the HRA.

### HRA vs. HRA KG

The HRA KG makes it possible to access HRA data efficiently and to ask biological questions via programmatic queries. Researchers can leverage the KG to enrich their assay data for deeper insights, e.g., supporting multiplexed antibody-based imaging with antibody panels, while clinicians can use it to explore biological questions, such as identifying the types and populations of epithelial cells in the human eye. The HRA KG is composed of multiple named graphs, each focusing on a specific part of the atlas, such as biological structures or spatial references. The HRA *collection* is a collection of HRA DOs with DOIs (ASCT+B Tables, 3D Reference Objects, OMAPs, etc.) that make up the core of the HRA at each release. When processed, it compiles to a (large) RDF graph and is hosted by the HRA KG at https://purl.humanatlas.io/collection/hra.

The HRA uses a KG (as opposed to a relational database)^31^ to ensure **(1) Flexibility**. The schema of the HRA KG can be extended as needed when new organs or HRA DO data types become available (as opposed to a rigid table schema that would need to be chosen for a relational database). Existing DOs can easily be updated. In a relational database, one would need a set of new tables. Many HRA DOs, such as 3D Reference Objects, are non-tabular and highly nested, which is challenging to model in relational databases. **(2) Support for disparate data**. HRA DOs take on many forms. For example, ASCT+B tables capture AS, CT in those structures, and the B that characterize them; they are linked to OMAP and Antibody Validation Report (AVR)^16^ tables (tabular data), 2D images and 3D models (graphic assets), as well as dataset graphs, such as scientific literature connected to the HRA (HRAlit)^32^ and cell type populations of the HRA (HRApop)^1^, all nested. A relational database would make it necessary to choose a schema for each of these DO types. A KG enables integration of many different DO types. **(3) Answering biological questions across HRA DO types**. The KG structure makes it possible to programmatically answer questions across multiple DOs for one entire organ via graph queries (e.g., the ASCT+B table for the kidney and the 3D Reference Object for the female, left kidney). In a relational database, this would necessitate a set of new tables that would need to be carefully created with foreign keys and relationships to support the kind of dynamic graph-based queries readily available in SPARQL. **(4) Deployment as 5-Star LOD**. RDF graphs can be imported into triple stores in their native format and easily be queried together with connected biomedical ontologies (genes, proteins, cells, anatomy), which are also published as RDF. Existing HRA KG queries bridge Uberon^7^, CL^19^, Provisional Cell Ontology (PCL)^33^, HUGO Gene Nomenclature Committee (HGNC, https://www.genenames.org)^34^, and HRA nodes, properties, and relationships stored or imported from their respective graphs.

To use the HRA KG in connection with a relational database, an initial query into the HRA KG can be used to retrieve data as a simple table; then, a database management system like PostgreSQL (https://www.postgresql.org/) can be used to aggregate the data with SQL features that perform complex window functions and aggregations on it. This way, data from the HRA KG can be indexed and used in a database downstream.

### Limitations

The current HRA KG has a number of known limitations that will be addressed in future HRA releases:

#### Automation

While many parts of the HRA KG construction process are automated, collecting and providing DOs in their original form, such as comma-separated values (CSV), binary Graphics Library Transmission Format (GLB, https://www.khronos.org/gltf), or scalable vector graphics (SVG), is still a manual process involving human labor. In future releases, we aim to employ machine learning algorithms^4,35–43^ to speed up tissue data segmentation and annotation, using human expertise to review (not hand-compile) DOs.

#### Build time

At present, building the HRA KG from unprocessed DOs using code in https://github.com/hubmapconsortium/hra-kg takes about 13 hours on a local Linux server with 256GB RAM and 20 cores. As new DO types are added and HRApop as well as HRAlit grow, we will continue to optimize the construction process by implementing better KG structures, using parallelization, and improving normalization and enrichment code (some libraries are particularly slow for certain DO types). Preliminary results from one experiment showed a nearly one-third reduction in execution time, demonstrating the potential of parallelization in faster, more efficient KG construction.

#### Reduce *ASCTB-TEMP* terms

As of HRA v2.2, 221 CT across 33 ASCT+B tables do not yet exist in CL or PCL; instead, they have an *ASCTB-TEMP* expert provided label. GitHub issues have been submitted for all, and the EMBL-EBI team is adding these terms. The **Technical Validation** section details ongoing efforts to link the HRA KG to existing ontologies. Additionally, as of February 2025, a total of 150 CT were added to CL^19^, and 468 CT were added to PCL^33^ by ASCT+B table editors. Additionally, 126 AS terms were added to the Uberon^7^ ontology.

##### Data modeling

To be most useful to the HRA KG, each new DO type must have a LinkML schema (see **Box 1**), normalization code, and enrichment code to transform the raw data into useful, queryable information. As new use cases are identified, the HRA structure and canned queries will be revised and expanded. Currently, HRAlit^32^ is being served via a relational database. Knowledge modeling is underway to create an HRAlit graph and to properly connect it to the HRA KG, which will allow users to query peer-reviewed literature and funded awards for entities in the HRA KG.

##### Ease of use

Retrieving data from KGs requires experience writing SPARQL queries, which few clinicians and biomedical researchers possess. The HRA KG comes with canned queries at https://apps.humanatlas.io/api/grlc/ as well as Jupyter Notebooks (see companion website at https://cns-iu.github.io/hra-kg-supporting-information). Going forward, we will create an HRA Developer Portal to help train and provide resources to users learning how to use the HRA KG. Additionally, since KGs offer access to structured data, we are exploring the possibilities of utilizing large language models (LLMs) to allow users to ask questions in prose. An LLM, enhanced by retrieval-augmented generation (RAG), was prototyped to support natural language queries that are informed by the knowledge in the HRA KG, see companion website at https://cns-iu.github.io/hra-kg-supporting-information/#using-llms-and-rag-with-hra-kg. Finally, we are working on an HRA KG Explorer UI, which will allow users to browse the KG via the web to quickly identify, select, and download HRA DOs of interest in all available graph formats. This will enable easy access to the HRA KG to users without experience writing code, making API requests via the https://grlc.io service, or running SPARQL queries.

## Methods

### HRA Digital Objects

HRA DOs come in diverse formats, such as ASCT+B tables^10^ (https://humanatlas.io/asctb-tables), 3D Reference Objects (https://humanatlas.io/3d-reference-library, https://3d.nih.gov/collections/hra), and OMAPs^16^ (https://humanatlas.io/omap), see **Box 1**. Each DO has a type, name, and version in a PURL (see **Box 1**). For example, the PURL for the ASCT+B table for the kidney is https://lod.humanatlas.io/asct-b/kidney/v1.6, where *asct-b/kidney/v1.6* indicates *asct-b* as the type, *kidney* as the name, and *v1.6* as the version. These DOs are provided by SMEs (see **Box 1**), then reviewed and validated throughout the HRA KG construction process.

HRA DOs, including ASCT+B tables, are well-defined data structures (see SOPs at https://humanatlas.io/standard-operating-procedures for details) that are regularly validated^44^, see also Technical Validation section. ASCT+B tables make it possible for anatomists, surgeons, and other experts to digitize knowledge about the CT and B in healthy tissue via Google Sheets. When constructing a table, SMEs are asked to crosswalk AS, CT, and B terms to ontology terms so they align with the HRA. Parsing of the Google Sheets is not advisable, as detailed validation and additional enrichment are required before the ASCT+B table data can be used in HRA construction or other workflows.

A complete list of the13 DO types in HRA v2.2 is provided in **Table 1**. These DOs can be categorized as reference data (*2d-ftu, asct-b, ctann, landmark, millitome, omap, ref-organ, vascular-geometry*), experiment data (*ds-graph, graph*), and other data (*collection, schema, vocab*). DOs are available in a variety of formats on the LOD server at https://lod.humanatlas.io. **Figure 1** illustrates high-level relationships among the 13 DO types.

**Figure 1.**
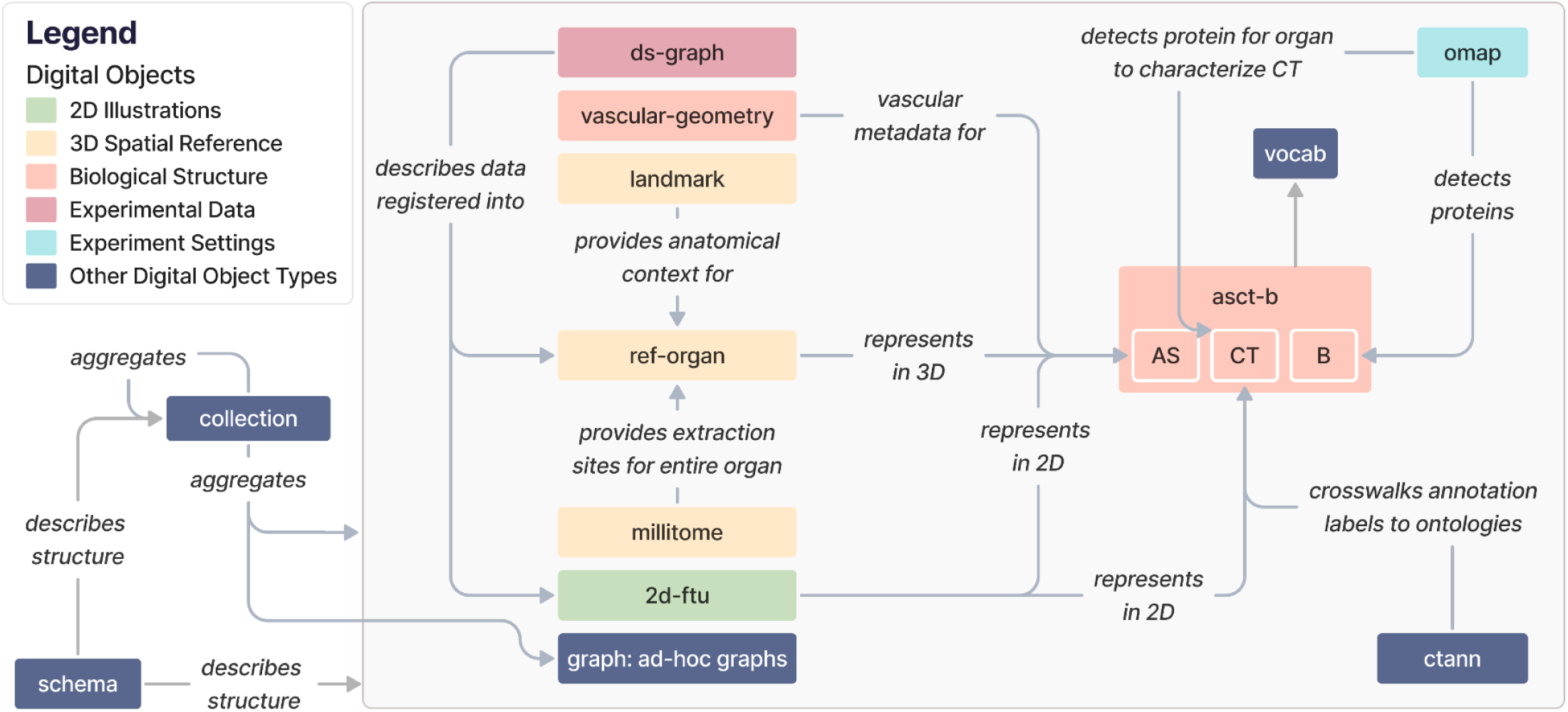
The 13 DO types in the HRA KG and how they relate to each other. Note that we replaced underscores in edge labels with blank spaces for legibility. Entity-relationship diagrams are provided on the companion website at https://cns-iu.github.io/hra-kg-supporting-information.

**Table 1.** Different DO types used in the HRA KG, describing their purposes and the data they contain plus SOPs detailing the construction of these DOs.

| Digital Object | Description |
| --- | --- |
| **Reference Data** |  |
| 2d-ftu | Provides 2D illustrations of FTU structures in an organ, with image assets and cell annotations that assign proper labels and identifiers based on CL for each image segment. An SOP is available<sup>45</sup>. |
| asct-b | Represents an ASCT+B table, see **Box 1**. Contains detailed knowledge about human body parts in a hierarchical order, explaining the organization of AS, the CT in each AS, and the B that distinguish each CT<sup>10</sup>. An SOP is available<sup>46</sup>. |
| ctann | Represents a crosswalk, see **Box 1**. Translates CT labels or abbreviations from sc/snRNAseq analysis tools, specifically Azimuth<sup>11</sup>, CellTypist<sup>12,13</sup>, and popV<sup>14</sup> into standardized terms in CL. The translation quality is measured using standard predicates, such as *exact match* and *narrow match* to ensure consistent data harmonization across sc/snRNAseq analyses. |
| landmark | Provides 3D model shapes representing features near organs of interest (e.g., an artery or pelvis bone near a kidney) to help experts accurately orient themselves when registering tissue blocks into a 3D Reference Object. |
| millitome | Provides data about cutting tissue samples using a millitome device. An SOP is available<sup>47</sup>. |
| omap | Represents an OMAP, see **Box 1**. Reduces the costs of conducting cell imaging experiments. OMAPs<sup>16</sup> contain a panel of antibodies designed to target specific proteins for identifying CT, AS, cell states, or cell membrane staining within organs, based on actual experimental projects. An SOP is available<sup>48</sup>. |
| ref-organ | Represents a 3D Reference Object, see **Box 1**. Provides 3D models of human organ structures, complete with accurate size and position data, to support the creation of a comprehensive 3D model of the human body, with each 3D model object carefully annotated with a proper label and an identifier from the Uberon and FMA ontologies. Multiple SOPs are available<sup>49,50</sup>. |
| vascular-geometry | Provides detailed geometry information on the human blood vascular system which captures key attributes of different vessels such as diameter and length, population, sample size, and reference to the source of data. Multiple SOPs are available<sup>51–54</sup>. |
| **Experiment Data** |  |
| ds-graph | Provides sample registration information submitted by consortium members in HuBMAP or other efforts, including accurate sample sizes and positions. When combined with *ref-organ* data, this information helps create 3D visual tissue sample placements. Additionally, the sample information is linked to datasets from researchers' assay analyses that offer deeper insights into the tissue samples. The “ds” stands for “dataset.” |
| graph | Contains externally created RDF (see **Box 1**) graph data that are useful for HRA use cases. |
| **Other Data** |  |
| collection | Combines multiple DOs to create a collection of data. The HRA itself is defined in the HRA KG as a curated *collection* of DOs in each release version. |
| schema | Describes the structure, i.e., the schema, of the normalized form of a single DO type, its metadata, or shared concepts between DOs. |
| vocab | Contains various reference ontologies and vocabularies that hold standard concepts and relationships used to construct the DOs. These DOs are typically external biomedical ontologies like CL and Uberon and provide a convenient mechanism for querying reference ontologies alongside HRA-curated DOs. |

The high-level relationships between DO types are as follows:

2D Illustrations (green): *2d-ftu* DOs illustrate AS (because FTUs are AS) and the CT in them, based on experimental data. *2d-ftu* DOs can be downloaded in their processed form or as SVG, Portable Network Graphics (PNG), or Adobe Illustrator (AI) files.

3D Spatial Reference (yellow): *ref-organ* DOs get anatomical context from *landmark* DOs, and *millitome* DOs provide extraction sites for an entire organ that a *ref-organ* DO represents. *ref-organ* and *landmark* DOs can be downloaded as GLB files. *millitome* DOs can be downloaded in the JavaScript Object Notation for Linked Data format (JSON-LD, https://json-ld.org).

Biological Structure (pink): The *asct-b* DO type plays a central role for multiple other DO types, see **Box 1** and **Table 1**. *vascular-geometry* DOs provide vascular metadata for *asct-b* DOs. Both can be downloaded in their raw distributions as CSV files.

Experimental Data (purple): *ds-graph* DOs describe experimental datasets mapped into a *ref-organ*. They can be downloaded as JSON-LD files.

Experiment Settings (cyan): *omap* DOs enable detection of proteins and CT and are thus connected to AS and B in *asct-b* DOs. *omap* DOs are specific to organs and assay types and can be downloaded as CSV files or Microsoft Excel Open XML Spreadsheets (XLSX).

*Other DO types (blue): ctann* DOs represent CTann crosswalks that map manual and machine learning annotations for CT from different CTann tools such as Azimuth^11^, CellTypist^12,13^, and popV^14^ to the ASCT+B tables via ontologies such as Uberon^7^, FMA^8,9^, CL^19^, PCL^33^. *ctann* DOs, like the aforementioned *omap* DOs, allow mapping experimental datasets, represented as the aforementioned *ds-graphs*, into the HRA. Both can be downloaded as CSV and XLSX files. *vocab* DOs are referenced by *asct-b*, *omap*, *ctann*, and *ref-organ* DOs to annotate AS, CT, and B with ontology terms and can be downloaded as Web Ontology Language files (OWL, https://www.w3.org/OWL). *graph* DOs are ad-hoc graphs that can reference any other DO type as needed, depending on their function and scope, and can thus have any download format available for the referenced DOs. All current *graph* DOs in the HRA KG are listed in **Supplemental Table 1.** *collection* DOs aggregate multiple DOs and can be downloaded as YAML files, allowing end users to create customized configurations for their particular needs. Importantly, the HRA itself is a collection DO, with the most recent release always available at https://lod.humanatlas.io/collection/hra/latest. All current collections in the HRA KG are listed in **Supplemental Table 2.** Finally, the *schema* DO type describes the structure of all HRA DO types plus their metadata and can be downloaded in a variety of formats, including YAML, PNG, and SVG (as an entity relationship diagram).

### Metagraph

HRA DO types can be aggregated into five thematic subgraphs: 2D Illustrations, 3D Spatial Reference, Biological Structure, Experimental Data, and Experiment Settings. The HRA KG metagraph in **Figure 2** depicts the higher-order relationships among these interconnected subgraphs. The 3D Spatial Reference subgraph (yellow) specifically is presented and explored in detail in a prior publication on the CCF Ontology^15^.

**Figure 2.**
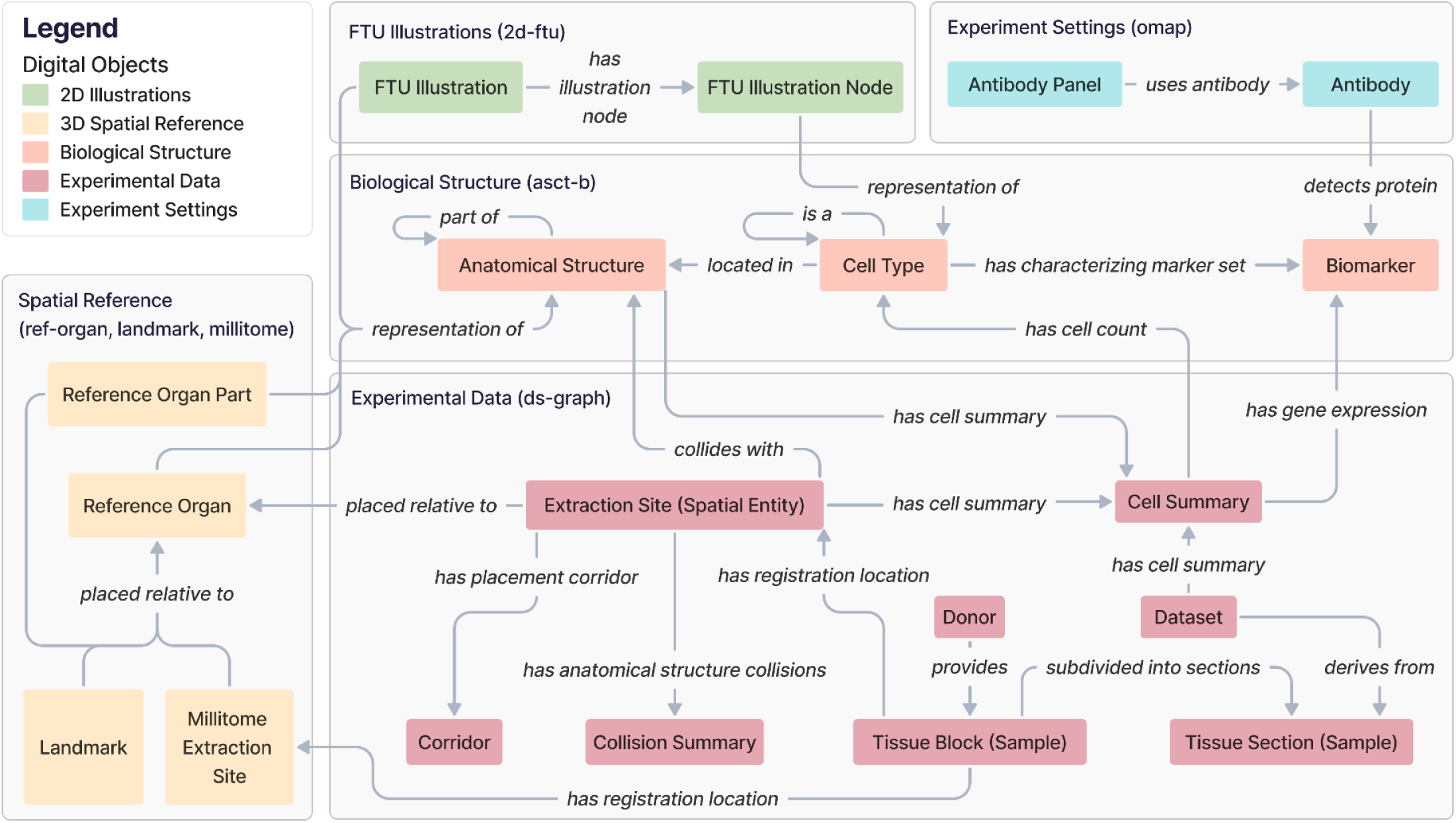
The HRA KG metagraph illustrates high-level relationships between subgraphs. Note that all edge labels have been modified as needed to avoid overlap while maintaining correct semantics. For class names (e.g., FTU Illustration), we added a blank space between CamelCased class names.

The **2D Illustrations** subgraph uses the Biological Structure subgraph to enrich its 2D FTU Illustrations and the FTU Illustration Nodes inside of it with the ontology-aligned naming for FTUs and CT. It does so via the Biological Structure subgraph to provide the 2D anatomical context for the AS and CT in the FTUs.

The **3D Spatial Reference** subgraph represents the 3D CCF used to accurately position 3D Reference Objects for organs, anatomical landmarks inside or adjacent to them, and millitomes (see **Table 1**) within the human body. This subgraph links to the Biological Structure to provide the 3D anatomical context for the AS.

The **Biological Structure** subgraph anchors all other components. It contains the AS, CT, and B from 33 ASCT+B tables together with their ontological relationships (for 32 organs plus one for anatomical systems). AS and CT can have self-loops, where an AS can be *part_of* another AS, creating a partonomy, and a CT can be a subclass of another CT in a typology (*is_a*).

The **Experimental Data** subgraph focuses on experimental Datasets generated from assay analyses performed on Donor Tissue Blocks (Samples). These are assigned an Extraction Site (Spatial Entity) with the HRA Registration User Interface (RUI)^6^ based on their anatomical origins to provide a location within the CCF. Since HRA v1.2, extraction sites are *placed_relative_to* Reference Organs; note that this is also a change in terminology, which used to be called *has_placement*, see prior publication^15^. All possible alternative locations of an Extraction Site (Spatial Entity), given its intersection(s) with one or multiple 3D AS, are captured in a Corridor. Systematic whole-organ registration is available as a Millitome, which defines a set of connected extraction sites *placed_relative_to* a Reference Organ. This subgraph also accommodates derived data computed from the assay results and extraction sites, such as (a) Cell Summaries, which provide cell type populations and mean gene expression values for specific CT and their associated datasets and 3D extraction sites, and (b) Collision Summaries, which identify AS that overlap with the registered tissue blocks inside a 3D extraction site, and detail the precise intersection volume and percentages in these collisions.

External annotations that are not shown in this subgraph are also possible: Datasets can be annotated with a publication; Donors, Tissue Blocks (Samples), and Datasets can have separate links to a data portal; and Donors can be annotated with a tissue provider.

Finally, the **Experimental Settings** subgraph catalogs Antibody Panels via OMAPs^16^, capturing details of specific Antibodies used to detect particular protein Biomarkers in the Biological Structure subgraph.

### The HRA KG in the HRA Ecosystem

The HRA KG represents major DOs of the HRA v2.2, including 33 ASCT+B tables, 23 OMAPs, 22 2D FTUs, 71 3D Reference Objects (plus two whole body models with all organs for male/female and a crosswalk from 3D AS to ontology terms), see https://apps.humanatlas.io/dashboard/data. In February 2025, the HRA KG had 10,064,033 nodes, 171,250,177 edges, and a size of 125.84 GB. The size of the 71 3D Reference Objects (GLB files, https://lod.humanatlas.io/ref-organ) in HRA v2.2 is 301 MB. In addition, the data covers anatomical landmarks which are used in the RUI to facilitate tissue block placement in 3D Reference Objects; these are available at https://lod.humanatlas.io/landmark. As of HRA v2.2, there are landmarks for 59 out of 71 3D Reference Objects. Together, they are 261 MBs large.

To make raw as well as processed HRA DOs available in a programmatic manner as RDF graphs, the HRA KG sits at the center of the HRA data ecosystem and serves as the primary database for the HRA, see **Figure 3**.

**Figure 3.**
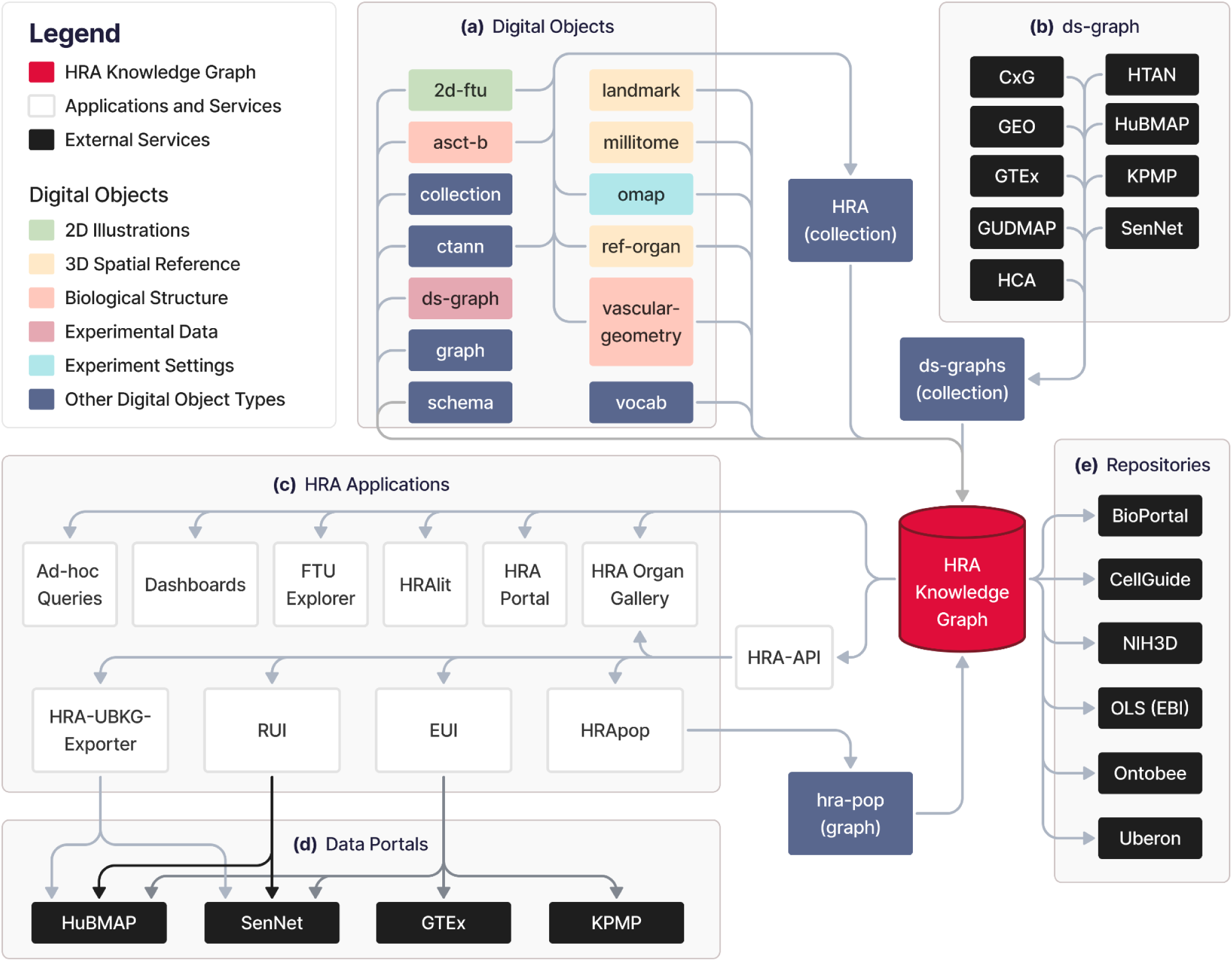
(a) HRA DOs, (b) experimental datasets, (c) HRA applications, (d) data portals, and (e) external services form the ecosystem around the HRA KG. Note that the HRA KG is able to serve all existing versions of the HRA and HRA DOs, incl. those preceding the most recent HRA v2.2 release in December 2024.

**(A)** 703 individual DOs of 13 types go through a 3-step process of normalization, enrichment, and deployment via the hra-do-processor (https://github.com/hubmapconsortium/hra-do-processor), where they are transformed from raw data in miscellaneous file formats into an RDF graph. Thus, the HRA KG integrates the knowledge from these DOs. Together, the HRA DOs form the HRA collection, which is the graph representation of the HRA. The normalization, enrichment, and deployment processes are described in the HRA KG Construction and Deployment section below.

**(B)** Graph representations of external experimental datasets from various sources with mean B expression values and cell type population data, resulting in the *ds-graph* DO type. Experimental data from various portals is mapped into the HRA via one or a combination of multiple methods, such as 3D tissue registration^6^, Azimuth^11^, CellTypist^12,13^, and popV^14^ annotations aligned to ASCT+B tables via ontology crosswalks, OMAPs^16^ for spatial proteomic data. These portals include the CZI CxG portal (https://cellxgene.cziscience.com), the Gene Expression Omnibus (GEO, https://www.ncbi.nlm.nih.gov/geo), the Genotype-Tissue Expression Portal (GTEx, https://gtexportal.org/home)^55^, the GenitoUrinary Developmental Molecular Anatomy Project (GUDMAP, https://www.atlas-d2k.org/gudmap)^56^, the Human Cell Atlas (HCA, https://www.humancellatlas.org/data-portal)^57^, the Human Tumor Atlas Network (HTAN, https://humantumoratlas.org/explore)^58^, the HuBMAP^25,26^ Data Portal (https://portal.hubmapconsortium.org), the KPMP^21,22^ Kidney Tissue Atlas (https://atlas.kpmp.org), and the Cellular Senescence Network (SenNet, https://data.sennetconsortium.org/search)^59^.

**(C)** HRA applications and services use the HRA KG as their main data backend through the HRA API or via a SPARQL endpoint (https://lod.humanatlas.io/sparql).

The HRA API provides the HRA UIs with access to Uberon, FMA, CL, PCL, and HGNC IDs for AS, CT, and B as well as spatial entities for tissue blocks and organs^15^ for the RUI and Exploration User Interface (EUI)^6^. For example, when using the EUI to select AS, CT, and B terms (on the left side of the UI), counts are retrieved from those relationships that are curated from multiple graphs. Since all graphs are in the RDF format, it is feasible to query across multiple graphs seamlessly without modifying the source graphs. The primary API at https://apps.humanatlas.io/api has programming language-specific client libraries in JavaScript, TypeScript, Angular 17+, and Python 3.6+. These client libraries are published to common code package managers, including NPM (https://www.npmjs.com) and PyPI (https://pypi.org), and they wrap the API calls into simple function calls to use from code, making HRA data easy to use from software development environments. A full list of client libraries is available at https://humanatlas.io/api. A set of example Python Notebooks is provided at https://github.com/x-atlas-consortia/hra-api/tree/main/notebooks. Links to publicly accessible instances of UIs using the HRA KG on data portals are provided in **Supplemental Table 3**. Ad-hoc queries to retrieve counts and access DOs from the HRA KG are easily possible via the SPARQL endpoint. The HRA Dashboard (https://apps.humanatlas.io/dashboard) and the HRA Portal (https://humanatlas.io)^1^ provide usage and data statistics about the HRA by querying the HRA KG. The FTU Explorer (https://apps.humanatlas.io/ftu-explorer)^60^ accesses CL IDs for cells and HGNC IDs for B via the HRA KG. HRAlit data^32^ (https://github.com/x-atlas-consortia/hra-lit) connects 136 DOs from HRA v1.4 to 583,117 experts, 7,103,180 publications, 896,680 funded projects, and 1,816 experimental datasets. The HRA Organ Gallery in virtual reality (VR)^61^ utilizes the HRA KG to show predicted CT in tissue blocks in immersive, 3D space. HRApop provides CT and mean B expressions for experimental datasets mapped to the HRA.

**(D)** The HRA KG is used in several **data portals**, including some from which *ds-graph* DOs are being extracted: HuBMAP, SenNet, GTEx, and KPMP. For example, the HRA-UBKG Exporter (https://github.com/x-atlas-consortia/hra-ubkg-exporter) is used to make HRA data available for HuBMAP Data Portal services (https://portal.hubmapconsortium.org), such as Uberon^7^ aligned organ pages (e.g., https://portal.hubmapconsortium.org/organ/lung), and AS search and filter functionality. The RUI and EUI are used in various portals to serve consortium-specific needs of tissue providers.

**(E)** Several **external repositories** serve and/or use HRA KG data: NCBO Bioportal and Ontobee host the HRA CCF Ontology^15^ (see **Box 1**) at https://bioportal.bioontology.org/ontologies/CCF and https://ontobee.org/ontology/CCFO. The OLS by EMBL-EBI provides a collection of HRA DOs for validation at https://www.ebi.ac.uk/ols4/ontologies/hra. CellGuide (https://cellxgene.cziscience.com/cellguide) utilizes the ASCT+B tables to identify and present canonical B for CT to their users. Finally, the NIH3D platform by the National Institute of Allergy and Infectious Diseases (NIAID) hosts all 71 3D Reference Objects for organs in the HRA v2.2 (https://3d.nih.gov/collections), plus two United files with all organs combined. Additionally, the NIH BioArt platform serves 22 2D FTU illustrations at https://bioart.niaid.nih.gov/discover?collection=2.

### HRA KG Construction and Deployment

All data and code needed to construct, deploy, and use the HRA KG are available at https://github.com/hubmapconsortium/hra-kg. A full list of all data and code is available in **Supplemental Table 4**. The HRA KG is constructed twice a year, coinciding with the HRA release cycle^1^ (see release notes at https://humanatlas.io/overview-training-outreach#release-notes). The most essential code piece, the **hra-do-processor** (https://github.com/hubmapconsortium/hra-do-processor), is built around three main components: the schema, the data processing pipeline, and the web infrastructure. The following sections detail each of these components.

#### Schema

Well-defined data schemas are crucial for ensuring data consistency, interoperability, and validation in data management and analysis. The HRA KG uses LinkML^18^, a flexible and user-friendly schema language designed to create effective data models and validation tools to ensure input data adheres to a defined schema. Entity relationship diagrams of core HRA schemas explain the relationships between different HRA DOs, see examples on the companion website at https://cns-iu.github.io/hra-kg-supporting-information/#mermaid-diagrams. A complete listing of the LinkML schemas used in HR KG construction is available in **Supplemental Table 5**. In addition to structural constraints, LinkML supports the implementation of reference integrity to ensure that linked entities conform to external ontologies. For instance, the *binds_to* slot in the *Antibody* class was explicitly constrained to accept only HGNC codes, preventing invalid associations with non-protein entities. Furthermore, LinkML enables explicit mappings between classes or slots to standardized vocabularies and ontologies. For example, the same *binds_to* slot was mapped to the *binds_to* property in the CCF vocabulary.

#### Data Processing Pipeline

The 13 HRA DO types described above come from SMEs, who contribute their knowledge of anatomy, antibodies, pathology, and experimental procedures via disparate data sources. Manually curated and experimental datasets from diverse sources need to be mapped to the HRA and standard ontologies, normalized to a standard format (e.g., unification of term labels), and enriched (e.g., linked to existing ontologies in support of causal reasoning). The **hra-do-processor** normalizes and enriches these DOs, then deploys them as RDF graphs. A catalog of these graphs is available on the HRA KG LOD server at https://lod.humanatlas.io. A sequence of five steps to convert different raw datasets into the HRA KG: normalization, enrichment, deployment, finalization, and serving. Implementation details are below.

##### Normalization

This initial step ensures that all incoming data is transformed into a consistent format that aligns with the predefined schema. The hra-do-processor loads and parses the disparate source data and transforms it into a standardized linked-data representation. For example, in the case of *asct-b* DOs, the source data comes from Google Sheets exported as CSV files (unnormalized, raw ASCT+B tables are available at https://humanatlas.io/asctb-tables); during normalization, the hra-do-processor reads this tabular structure and converts it into the tree structure shown in **Supplemental Figure 1.** An exemplary YAML file is provided at https://github.com/cns-iu/hra-kg-supporting-information/blob/main/docs/intermediary_format.yaml. YAML was chosen as the standard file format for the normalized data due to its simplicity, readability, and interoperability with JSON, and high level of support in LinkML. By converting the LinkML schema into a JSON schema file, we can easily validate the translated YAML data to ensure it adheres to the defined schema and our automated ingestion code is correctly implemented.

In general, HRA DOs come in two major formats: tabular data (like *asct-b* DOs) and nested key-value pairs. Extracting knowledge from tabular data begins with schema inference, which includes identifying header rows, data types, and relationships between columns. Typically, the first row serves as the header, and the data types are inferred by analyzing the entire column values. Determining the relationships between columns requires either consulting a knowledge base or relying on domain experts. In *asct-b* DOs, the relationship between the column headers for AS at different levels (*AS/1*, *AS/2*, *AS/3*) involves aligning their values with Uberon or FMA and confirming that they follow the *part_of* relationship. Further refinement means identifying any parent-child relationships by grouping columns that logically belong together as a single concept. For example, the columns *rrid*, *host*, *isotype*, *clonality*, *conjugate* in an *omap* DO are attributes of an antibody rather than separate concepts. Once the table structure is understood, the next step is semantic annotation, where table components are linked to concepts from external knowledge sources or ontologies. This process involves three key actions: (1) matching individual cell values to ontology terms, such as linking the label *Kidney* to *UBERON:0002113*; (2) associating entire columns with controlled vocabularies, for example, defining the *REF/1* column as *dct:references* to describe a cited, related resource, and (3) associating relationships between columns to ontology terms, such as linking the relation between *AS/2* and *AS/1* columns to *BFO:0000050*, which denotes a *part_of* relationship.

For nested key-value pairs data, schema inference is more straightforward since the hierarchical structure is more visible than in flat, tabular data. However, it is still necessary to determine data types, structural relationships, and references within the data. For instance, the *size* field in data from *ref-organ* DOs is recognized as a nested object that contains *x*, *y*, and *z* fields, which are later inferred to be of decimal type based on the populated values. When identifying references, fields containing the substring *id* are typically recognized as reference fields that indicate a link to an external entity.

Once the schema is inferred, a similar step of semantic annotation is performed to the data elements by linking them to ontology concepts. For example, a nested field *node.size.x* is mapped to the *x_dimension* annotation property in the CCF Ontology. Likewise, the value *VH_M_aortic_valve* is mapped to *UBERON:0002137*, allowing it to be correctly identified as an aortic valve from Uberon, complete with its anatomical context. Finally, references are introduced using unique identifiers to maintain consistency and avoid redundancy. For instance, Research Resource Identifiers (RRID, https://www.rrids.org) are used to refer to a specific antibody rather than repeating its full details.

#### Enrichment

This step converts the normalized data to RDF and enriches it with relationships, entities, and metadata from external resources. After the source data is translated into YAML and validated, enrichment begins by converting the validated data into OWL-based statements. The LinkML framework offers tools that facilitate the transformation of tree-structured data into OWL constructs, including class and property declarations, as well as instances of a class (see **Supplemental Figure 2**). OWL was chosen as the data representation for the enrichment step due to its robust capabilities for rich data expression, its ability to embed semantic meaning, and its seamless integration with LinkML. In turn, LinkML provides direct support for OWL by allowing schema elements to include OWL constructs, making it easy to map the data into a semantically rich ontology structure. The enrichment process continues by integrating additional information from reference ontologies as well as authoritative databases like RRID and the Antibody Registry API (https://www.antibodyregistry.org) to retrieve metadata (label, description) about antibodies, for which there is only an RRID in the raw HRA DO for OMAPs.

The goal is to enhance the initial data gathered from SMEs with more detailed, authoritative information. In *asct-b* DOs, many data points already reference Uberon and CL terms. We enrich these terms by retrieving supplementary information from the corresponding ontologies, including class hierarchies, labels, definitions, synonyms, database references, and visual depictions. For example, we identified the standard label for a *CL:0002306* as *epithelial cell of proximal tubule* which is categorized under the broader class *meso-epithelial cell*. These details, which were absent from the original dataset, add valuable context. The end result is a semantically enriched dataset that not only preserves the original data but also extends it with additional context, relationships, and meaning.

##### Deployment

Once the data is enriched, it is prepared for use in downstream applications or for access by end users. This stage involves organizing the data into its final distribution formats and setting up the correct structure for the file system directory. HRA data is used by many different tools that need diverse formats: HRA UIs like the EUI and RUI^6^ use JSON-LD, which is best when using the data directly and imperatively (i.e., in a programming language using *for* loops). Python and JavaScript have native support for handling JSON and have semantics built in, and an exemplary Jupyter Notebook (https://jupyter.org) to showcase how an ASCT+B table as a JSON file can be parsed is provided on the companion website at https://cns-iu.github.io/hra-kg-supporting-information; the Usage Notes section also provides background. The Blazegraph (https://blazegraph.com) SPARQL server uses Terse RDF Triple Language (Turtle, https://www.w3.org/TR/rdf12-turtle); the Turtle format also helps developers write SPARQL queries to the HRA KG by making its triple structure explicit and showing possible subjects, predicates, and objects. Older semantic web tools use RDF/Extensible Markup Language (XML, https://www.w3.org/TR/rdf-syntax-grammar), N-Triples (https://www.w3.org/TR/n-triples/), and N-Quads (https://www.w3.org/TR/n-quads). Additionally, Robot (http://robot.obolibrary.org/convert.html), Apache Jena and RDF I/O technology (RIOT, https://jena.apache.org/documentation/io) use XML for reifying graphs. HRA KG data is preprocessed in those formats to be readily usable by others. Publishing all these formats streamlines the content negotiation process later (see **Box 1**) when different applications access the published HRA KG on the LOD server at https://lod.humanatlas.io, which can then immediately deliver the HRA data in the correct format. During the deployment step, the hra-do-processor also prepares the metadata that accompanies the graph data before copying files and data assets into their designated folders.

##### Finalization

Next, the necessary metadata and landing pages for web publication are generated, e.g., https://lod.humanatlas.io/asct-b/eye/latest leads to the most recently published ASCT+B table for the eye. In addition, this stage includes building the SPARQL database that will be uploaded to the web for users to access at https://lod.humanatlas.io/sparql. In the deployment step, data and metadata for each DO are converted and exported. Finalization derives additional files across all DOs, including metadata catalogs, latest versions of each DO, HTML landing pages to navigate the HRA KG and view DOs, and an indexed and optimized database file for the Blazegraph SPARQL server. The database contains the latest version of every DO, every version of the HRA collection, and a metadata catalog that contains metadata for every version of every DO in the HRA KG.

##### Serving

Data processed in the previous steps, including raw DO data, processed data products, HTML pages, metadata, and the SPARQL database, are made available online at https://lod.humanatlas.io. The data is regularly updated and synchronized, either during scheduled releases or when updates occur, to ensure that the most current version is always available. To make the processed data widely accessible, Amazon Web Services (AWS, https://aws.amazon.com) is used to serve the HRA KG as linked open data, employing three of its core services: S3, ECS and CloudFront for data storage, computation, and content delivery, respectively. Implementation details are provided in the next section.

#### Web Infrastructure

*Amazon S3 (Simple Storage Service)* is a highly scalable data storage service to store and retrieve data. The HRA KG uses S3 to store the content from the local deployment directories, including the Blazegraph database file. By syncing these local directories with an S3 storage, the data is securely stored and readily available for content delivery.

*Amazon ECS (Elastic Container Service)* is a fully managed container service to run applications in Docker containers for a highly scalable and reliable environment for our computation needs (https://www.docker.com). For HRA, a Blazegraph instance is run within an ECS container. The ECS container periodically checks the S3 storage for an updated Blazegraph database file. When a newly built Blazegraph file is detected, ECS will seamlessly update the Blazegraph server to ensure that the latest data is available for querying.

*Amazon CloudFront* is a global Content Delivery Network (CDN) designed to accelerate the distribution of content by caching copies at multiple serving locations around the world. The HRA KG uses CloudFront to create a URL fabric that caches and serves content from S3 storage to ensure fast and reliable access for users, regardless of their geographical location. The content stored in S3 is made publicly available through URLs leading to a CDN, e.g., https://cdn.humanatlas.io/digital-objects/ref-organ/liver-female/v1.2/assets/3d-vh-f-liver.glb, which returns the GLB file for the 3D Reference Organ for the female liver. Additionally, CloudFront provides advanced content negotiation features through Amazon CloudFront functions, enabling the dynamic handling of URLs starting with https://purl.humanatlas.io and https://lod.humanatlas.io. Content negotiation allows the web infrastructure to serve data in different formats based on user needs, whether a user requires RDF, JSON, XML, or another format. The PURL returns HRA DO *data* as an RDF graph based on the *Accept* header of the request: human users get redirected to the LOD server, machines to JSON or RDF versions. The LOD server also returns metadata for processed HRA DOs, such as who created it, when it was published, and what different assets and reifications are available to download, as Data Catalog Vocabulary (DCAT) datasets with provenance (https://www.w3.org/TR/vocab-dcat-3) based on the *Accept* header: Human users get HTML, machines get JSON or RDF. Moreover, CloudFront also acts as an intermediary for the SPARQL endpoint hosted by Blazegraph within ECS by making it accessible at https://lod.humanatlas.io/sparql.

### Other Ontologies

The HRA KG includes other reference ontologies at https://lod.humanatlas.io/vocab (e.g., Uberon and CL) so they can be queried together in an efficient manner. **Table 2** lists all ontologies that are included in the HRA KG together with their version numbers.

**Table 2.** Ontologies used in the HRA KG as of HRA v2.2.

| Name | Description | Version Number | URL | Main Website |
| --- | --- | --- | --- | --- |
| CCF | Common Coordinate Framework Ontology<sup>15</sup> | 3.0 | https://purl.humanatlas.io/vocab/ccf | https://humanatlas.io/ccf-ontology |
| CL | Cell Ontology<sup>19</sup> | 2024-09-26 | https://purl.humanatlas.io/vocab/cl | https://obophenotype.github.io/cell-ontology |
| FMA | Foundational Model of Anatomy<sup>8,9</sup> | 5.0.0 | https://purl.humanatlas.io/vocab/fma | http://si.washington.edu/projects/fma |
| HGNC | HUGO Gene Nomenclature Committee<sup>34</sup> | 2024-03-04 | https://purl.humanatlas.io/vocab/hgnc | https://www.genenames.org |
| HRAVS | HuBMAP Research Attributes Value Set | 2.5.3 | https://purl.humanatlas.io/vocab/hravs | https://bioportal.bioontology.org/ontologies/HRAVS |
| LMHA | Cell Ontology for Human Lung Maturation (LungMAP Human Anatomy)<sup>62</sup> | 1.4 | https://purl.humanatlas.io/vocab/lmha | https://bioportal.bioontology.org/ontologies/LUNGMAP_H_CELL |
| PCL | Provisional Cell Ontology<sup>33,63</sup> | 2024-07-11 | https://purl.humanatlas.io/vocab/pcl | https://github.com/obophenotype/provisional_cell_ontology |
| RO | OBO Relation Ontology (https://doi.org/10.5281/zenodo.32899) | 2024-04-24 | https://purl.humanatlas.io/vocab/ro | https://github.com/oborel/obo-relations |
| Uberon | Uberon Multi-species Anatomy Ontology<sup>7</sup> | 2024-11-25 | https://purl.humanatlas.io/vocab/uberon | https://obophenotype.github.io/uberon |
| VCCF | Vasculature Common Coordinate Framework<sup>10,64</sup> | 2024-02-23 | https://purl.humanatlas.io/vocab/vccf | https://github.com/hubmapconsortium/hra-vccf |

## Data Records

The whole HRA KG is available on https://cdn.humanatlas.io/hra-kg-releases/hra-kg.v2.2.tar.xz and is ∼5.1 GB large. The primary server for the HRA KG v2.2 is at https://lod.humanatlas.io. The SPARQL endpoint to query the HRA KG is at https://lod.humanatlas.io/sparql. The HRA API (https://apps.humanatlas.io/api) supports programmatic access to the HRA KG. Exemplary queries are available via the companion website at https://cns-iu.github.io/hra-kg-supporting-information.

BioPortal hosts (1) the HRA at https://bioportal.bioontology.org/ontologies/HRA (mirror of https://purl.humanatlas.io/collection/hra/v2.2) and (2) the CCF Ontology at https://bioportal.bioontology.org/ontologies/CCF (mirror of https://lod.humanatlas.io/vocab/ccf).

OLS hosts the latest versions of both the HRA and CCF at https://www.ebi.ac.uk/ols4/ontologies/hra and https://www.ebi.ac.uk/ols4/ontologies/ccf, respectively. OLS provides both a web-based GUI for users and programmatic access via the OLS REST API (https://www.ebi.ac.uk/ols4/help), enabling the HRA and CCF to be accessed using the same standard interface as other ontologies.

The NIH3D platform by NIAID hosts all 71 3D Reference Objects for organs in the HRA v2.2 alongside two United files with all organs combined (https://3d.nih.gov/collections). 22 2D FTU illustrations are at https://bioart.niaid.nih.gov/discover?collection=2.

Weekly run term and relationship validation reports of ASCT+B Tables are available at https://github.com/hubmapconsortium/ccf-validation-tools/tree/master/reports.

All data and SOPs are released under Creative Commons Attribution 4.0 International (CC BY 4.0).

## Technical Validation

HRA KG validation covers comparison to other knowledge graphs; growth of the HRA coverage and usage over time; and term additions to Uberon and CL based on HRA and other atlasing efforts.

### Comparison to other KGs

We compared the HRA KG with other major biomedical KGs quantitatively (number of nodes, edges, edge types, and size) and qualitatively (technology used, accessibility via SPARQL, need for authentication, presence of API for canned queries, license, and reproducibility), see **Table 3**. Jupyter Notebooks with queries for stats in support of this comparison is on the companion website at https://cns-iu.github.io/hra-kg-supporting-information#comparison-to-other-kgs.

**Table 3.** Key properties of HRA KG and other KGs.

| Property | HRA KG | SPARC KG | Petagraph | UBKG (Data Distillery) | Ubergraph | ORKG |
| --- | --- | --- | --- | --- | --- | --- |
| #Nodes | 1,543,074 | 188,146 | 32,192,544 | 54,154,737 | 4,071,817 | 557,821 |
| #Node types | 1,540,507 | 164,750 | 3 | 5 | 3,942,352 | 556,706 |
| #Edges | 101,720,865 | 812,969 | 151,690,690 | 227,826,739 | 458,605,464 | 1,853,694 |
| #Edge types | 509 | 221 | 1,960 | 2,260 | 778 | 4,217 |
| Size* | 30,848 MB | 789 MB | 138,571 MB | 162,895 MB | 96,450 MB | 437 MB |
| Date of latest release | Dec 15, 2024 | Sept 21, 2024 | May 5, 2024 | Jan 3, 2025 | Jan 13, 2025 | Feb 11, 2025 |
| Technology | RDF/ SPARQL | RDF/ SPARQL | Neo4J | Neo4J | RDF/ SPARQL | RDF/SPARQL (Neo4J also available) |
| Provides SPARQL endpoint | X | X |  |  | X | X |
| No authenticated access needed for queries | X | X |  |  | X | X |
| Has API with canned queries | X | X | X | X | X | X |
| Data Usage License (strictest listed) | CC BY 4.0 | CC BY 4.0 | UMLS License | UMLS License | CC BY 4.0 | CC0 1.0 Universal** |
| Data and code full reproducible | X | X | X | X | X | X |

\*RDF-based KGs were converted to N-Quads to get the total uncompressed size for each. To compare sizes, Neo4J KGs were converted to RDF N-Quad format; context and documentation are at https://cns-iu.github.io/hra-kg-supporting-information/#comparison-to-other-kgs. A Neo4J utility (https://neo4j.com/labs/apoc/4.1/export/json) was used to export a whole database in the JSON-lines format (https://jsonlines.org) and converted to RDF using a JSON-LD context with newline-delimited JSON (ndjsonld, https://www.npmjs.com/package/ndjsonld).
\*\*Data sourced from Papers With Code, licensed CC BY-SA.

The Stimulating Peripheral Activity to Relieve Conditions (SPARC) KG (https://sparc.science/tools-and-resources/6eg3VpJbwQR4B84CjrvmyD)^65^ provides FAIR vocabulary for its multimodal models, data, maps, and simulations across species. It ingests community ontologies across SPARC-relevant domains, such as physiology, anatomy, molecular structures, and experimental design (see **Figure 4**) and serves the data via an endpoint at https://blazegraph.scicrunch.io/blazegraph/sparql. Petagraph^24^ and UBKG were introduced in the Background & Summary section. Petagraph and UBKG data were retrieved from the Common Fund Data Ecosystem (CFDE) Data Distillery (https://dd-kg-ui.cfde.cloud/about). Ubergraph (https://github.com/INCATools/ubergraph)^66^ is a RDF triplestore with a public SPARQL endpoint that makes 39 ontologies from the Open Biological and Biomedical Ontologies (OBO) Foundry (http://obofoundry.org) available as a pre-computed knowledge graph to support ontology browsing and connection verification. Finally, the Open Research Knowledge Graph (ORKG, https://orkg.org)^67,68^ aims to improve processing of scholarly knowledge via an infrastructure that makes the description of research contributions machine-readable; it uses large language models to support natural language queries (https://ask.orkg.org).

**Figure 4.**
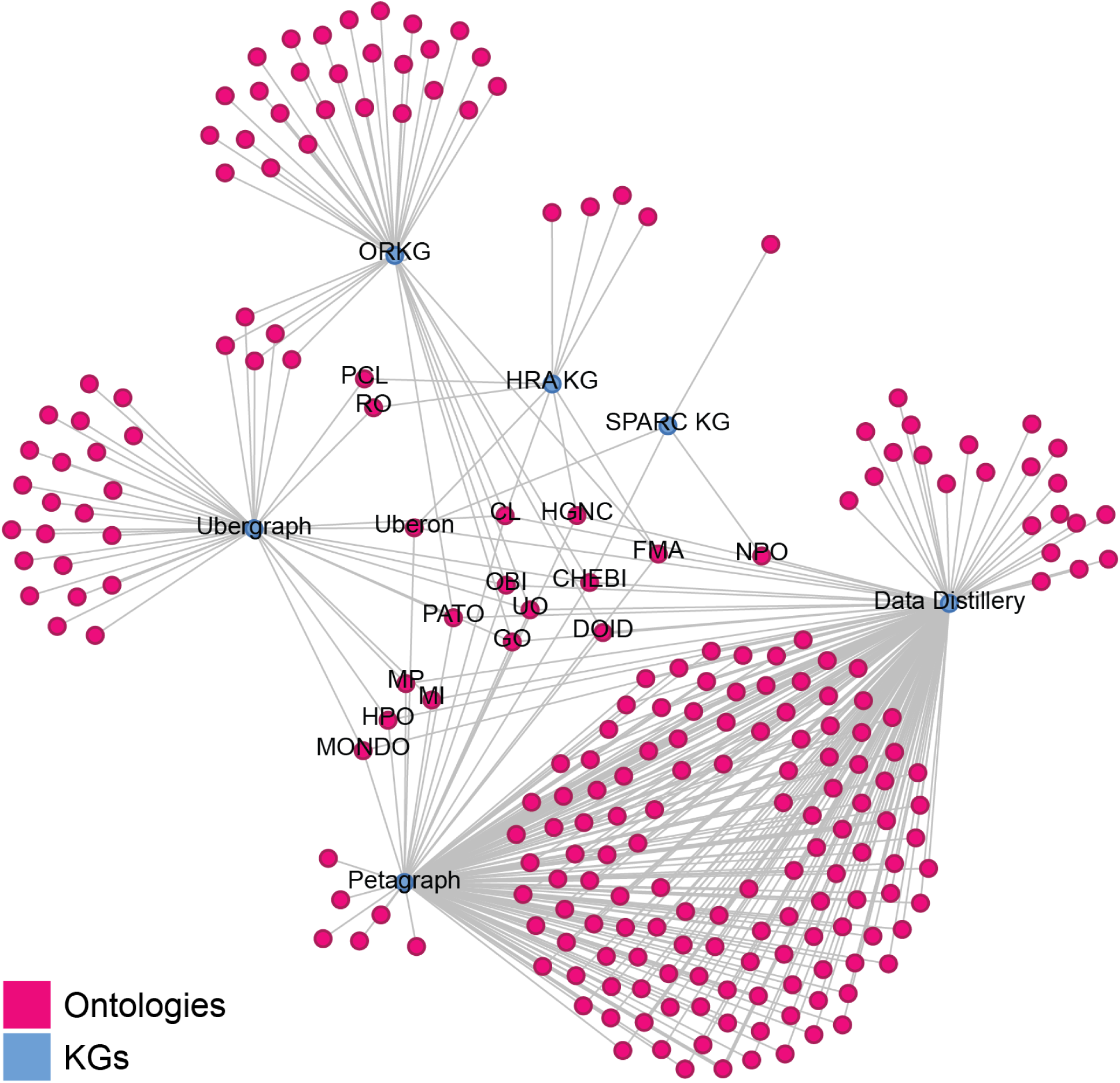
Bimodal network of KGs and the ontologies they ingest and serve. Automatic labels are only shown for nodes with a degree greater than or equal to three, i.e., the KGs themselves and all ontologies shared by at least three KGs. Some ontologies have manually added labels (PCL, RO). Note that Petagraph and Data Distillery ingest large amounts of ontologies that are not shared between KGs and are thus very similar. The most shared ontologies are Uberon, CL, FMA, HGNC, CHEBI, STRING, DOID, UO, PATO, OBI, and GO. The layout was made with Gephi (https://gephi.org), using the Yifan Hu (Proportional) Algorithm for layout (http://yifanhu.net/PUB/graph_draw.pdf).

The number of nodes and complexity of these KGs ranges from 188k nodes of 165k types to 54M nodes of 5 types. Size ranges from 437 MB for ORKG to 162GB for UBKG (Data Distillery version).

The UBKG and Petagraph use the Neo4J platform and require a UMLS License (https://www.nlm.nih.gov/databases/umls.html). ORKG has a CC0 1.0 Universal license. All other KGs are open access with the CC BY 4.0 license (https://creativecommons.org/licenses/by/4.0/deed.en).

**Note**: One complication when comparing KGs is that data representation is different between RDF-based KGs using RDF/SPARQL and Property Graph (PG)-based KGs using Neo4J. RDF consists of only edges (triples) with subject, predicate, and object. PGs consist of a set of nodes and edges. In RDF, nodes (i.e., resources) are annotated via an edge/triple (e.g., <https://example.com> <http://www.w3.org/2000/01/rdf-schema#label> “Example.com Label”), whereas in a PG, annotations are just part of the node’s data structure. So, in RDF, edges are used to both annotate nodes and represent relationships between nodes. As a result, comparing raw edge counts between a PG and RDF graph is complicated. To account for this, we wrote corresponding SPARQL (for RDF Graphs) and Cypher (for PGs, https://neo4j.com/docs/cypher-manual/current/introduction) queries to retrieve the number of nodes, edges, and edge types that, while not perfect, still gives a better sense of how their quantitative relationship. A full list of SPARQL and Cypher queries can be found in the Supporting Information at https://cns-iu.github.io/hra-kg-supporting-information/#comparison-to-other-kgs.

Figure 4 shows the bimodal network of the seven KGs (blue nodes) and the 288 ontologies (pink nodes) they import and serve. All but ORKG import Uberon and CL and as a result, they cover the same organs, anatomical structures and cell types that exist in these ontologies. The HRA KG has 951 additional TEMP-ASs, 221 TEMP-CTs, and 296 TEMP-Biomarkers (genes, proteins, lipids, metabolites, proteoforms) that were identified by human experts as missing and will be added to Uberon, CL, and biomarker ontologies (https://api.triplydb.com/s/e771any9d).

### Growth of Uberon, CL, and PCL over time from other sources

To compare the growth of the HRA KG against other KGs, the number of nodes, edges, and edge types was computed for Uberon, CL, and PCL. To that end, OWL files with release dates were downloaded from GitHub repositories (Uberon: https://github.com/obophenotype/uberon/releases, CL: https://github.com/obophenotype/cell-ontology/releases, PCL: https://github.com/obophenotype/provisional_cell_ontology/releases), then queried with a SPARQL query. The resulting CSV file holds the number of nodes, edges, and edge types for Uberon, CL, and PCL by release. All data and code is available in the Supporting Information at https://github.com/cns-iu/hra-kg-supporting-information/tree/main/notebooks. The CSV file at https://github.com/cns-iu/hra-kg-supporting-information/blob/main/notebooks/output/other-ontologies-growth.csv shows the growth of these three KGs between 2022 and 2025. Note that Uberon was first started in 2012, but the data available via releases in GitHub only goes back to 2022.The number of edge types is relatively stable, hovering around 100 for all three. The number of nodes and edges has been stable in Uberon and CL but has seen a steep increase for PCL since mid-2024, highlighting the more rapidly changing nature of PCL versus the more stable and established other two ontologies. Since 2022, Uberon/CL/PCL have had at most 15,959/3,122/16,980 nodes, 238,139/30,635/221,269 edges, and 171/58/85 edge types.

### Atlas Coverage

The HRA Dashboard (https://apps.humanatlas.io/dashboard/data) shows the size and coverage of HRA data; the number and type of HRA usage over time; publication and experimental data linked to the HRA; plus diversity of HRA authors, tissue donors, and users.

Figure 5a shows the number of instances of different DO types. Specifically, HRA KG v2.2 covers 71 organs via 3D *ref-organ* DOs (note that eye, Fallopian tube, kidney, knee, mammary gland, ovary, palatine tonsil, renal pelvis, and ureter) have left and right HRA DOs for the same Uberon ID and that exactly five organs are female only (fallopian tube, mammary gland, ovary, placenta, uterus) while one is male only (prostate). 33 *asct-b* DOs tables exist, where 32 cover organs and one covers anatomical systems. *ctann* crosswalks exist for 23 organs, with 12 organs having crosswalks for more than one CTann annotation tool. At present, *omap* DOs exist for 13 organs, covering different assay types. Finally, *2d-ftu* DOs are present for 10 organs, with the kidney having the most (eight).

**Figure 5.**
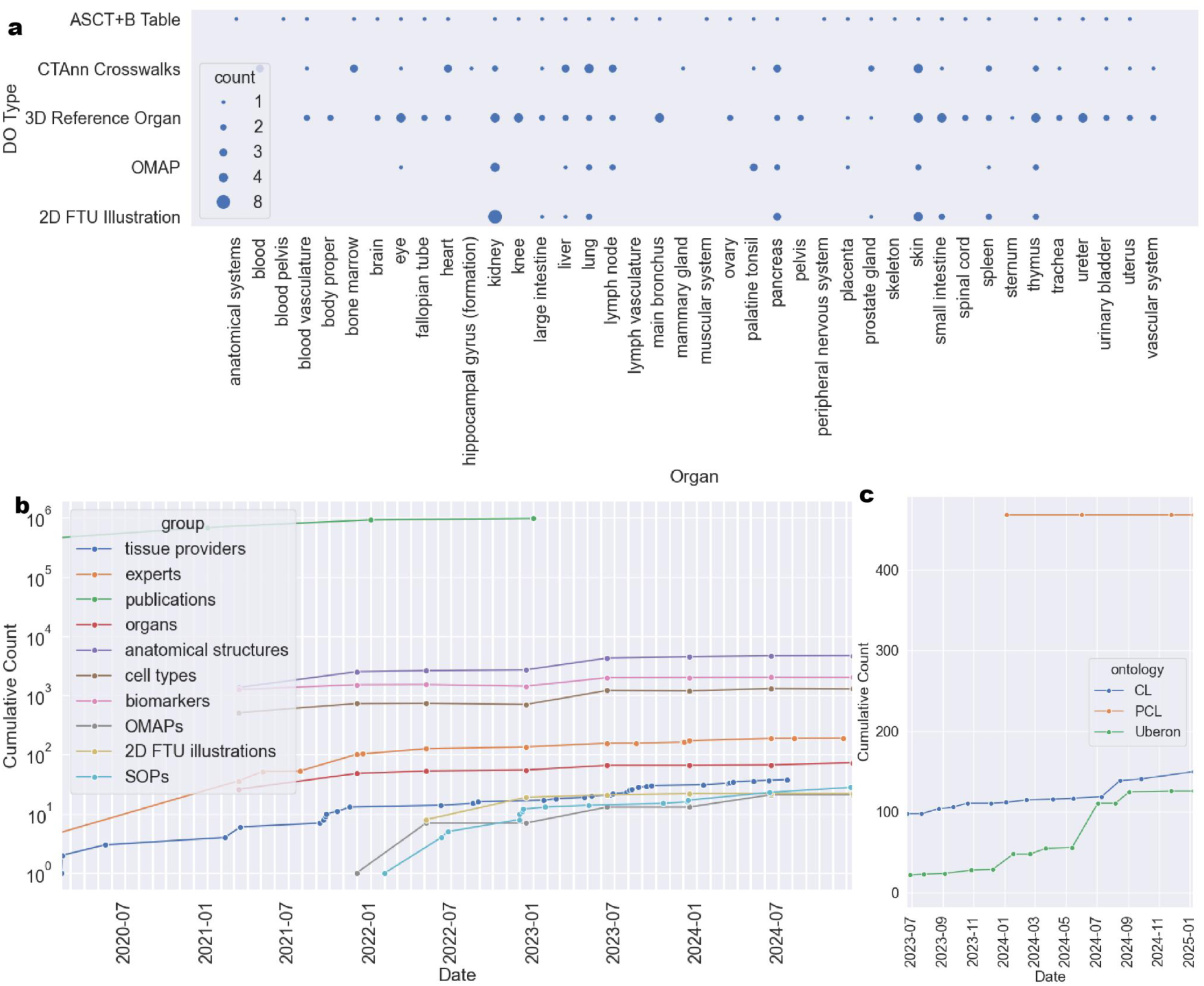
Graphs from the HRA Dashboard showing HRA KG v2.2 growth and coverage. a: DOs per organ. b: HRA KG growth in terms of DOs over time. c: Number of terms added from the HRA to CL, PCL, and Uberon over time.

Figure 5b plots HRA growth since HuBMAP started in 2018 (but only shows data since March 2020). The green line shows publications linked to the HRA via the HRAlit^32^ dataset, which captures literature through early 2023 connected to the HRA v1.4 (June 2023). In 2021, more HRA DOs were added (*omap* DOs^16^, later *2d-ftu* DOs^60^) plus a growing library of SOPs detailing HRA construction and usage (https://humanatlas.io/standard-operating-procedures). As new HRA DOs are published, they are ingested into the HRA KG as it is constructed and deployed for usage.

Figure 5c shows the number of terms that were added to existing ontologies. Between 2018 and February 2025, the HRA and other atlasing efforts added 126 terms to Uberon, 150 to CL, and 468 to PCL (shown are additions since July 2023).

## Usage Notes

Users access the HRA KG via UIs, APIs, and data products on https://lod.humanatlas.io to answer biomedical questions. A list of all HRA applications that use the HRA KG is provided in **Supplemental Table 3.** A list of publications and aliases used throughout HRA applications per HRA DO is provided in **Supplemental Table 6.**

The HRA KG makes it possible to access harmonized, high-quality reference and experimental data in standard data formats. Three widely used queries are detailed in this section: **(1) retrieve AS-CT-B records** from the ASCT+B tables**, (2) get mean B expression values** for CT across datasets in HRApop, and **(3) query the HRA KG to achieve two types of prediction**s: cell type populations given a spatial origin (3D extraction site), and spatial origin (3D extraction site and registration corridor) given a cell type population, see HRA user stories 1-2 in a related publication^1^.

To simplify HRA KG usage, we use the https://grlc.io service to make a set of canned SPARQL queries available to execute as simple web requests. Internally, the service creates an OpenAPI specification (https://swagger.io/specification) that advertises the available queries. We provide a user-friendly interface to these queries at https://apps.humanatlas.io/api/grlc/. Annotated screenshots of the query interface with instructions on how to run the queries and download the resulting data is available on the companion website at https://cns-iu.github.io/hra-kg-supporting-information/#how-to-run-queries-via-our-openapi-spec. This deployment was inspired by the PubMed MeSH SPARQL Explorer at https://id.nlm.nih.gov/mesh/query.

### Get records from ASCT+B tables

The HRA KG API makes it easy to retrieve AS-CT-B records, properties, or counts for one or multiple organs.

**Get ASCT+B counts for all tables**: Retrieve the number of unique AS, CT, and B terms across all ASCT+B tables via this query for the latest HRA release:

- https://apps.humanatlas.io/api/grlc/hra.html#get-/as-ct-b-counts

**Returns**: A table with the number of unique AS, CT, B in the latest version of the HRA.

Retrieve the number of unique AS, CT, and B terms across all ASCT+B tables for all HRA releases:

- https://apps.humanatlas.io/api/grlc/hra.html#get-/as-ct-b-counts-all-versions

**Returns**: A table with the number of unique AS, CT, B across all versions of the HRA (not just the latest one). Note that the only difference between the two queries is the added *FROM HRA:* statement in the SPARQL query, which limits the SPARQL search pattern of the first query to the latest HRA collection at https://purl.humanatlas.io/collection/hra.

**Get ASCT+B records for one organ**: Given the PURL of an *asct-b* DO, retrieve ASCT-B records from the ASCT+B table:

- https://apps.humanatlas.io/api/grlc/hra.html#get-/asctb-in-table.

**Returns**: A table with one row per AS-CT-B record in the specified table, in the format: AS, CT, B labels, and the AS, CT, B ontology ID (if crosswalked, otherwise it returns a temporary *ASCTB-TEMP* ID). Exemplary Python code that uses this endpoint is provided on the companion website at https://cns-iu.github.io/hra-kg-supporting-information#basic-usage. The user can choose from common data formats (CSV or JSON) for the response via the *Accept* header.

**Get ASCT+B records for all organs:** Retrieve individual records for all ASCT+B tables (rather than just counts or records for one organ):

- https://github.com/x-atlas-consortia/hra-pop/blob/main/queries/hra/asctb-records.rq

**Returns**: A table with one row per AS-CT-B record for all ASCT+B tables, in the format: AS, CT, B labels, and the AS, CT, B ontology ID (if crosswalked, otherwise it returns a temporary *ASCTB-TEMP* ID). Because the result of this query is large (1,048,576 rows and a total of 325 MB) and takes longer to run, it has not been deployed via https://grlc.io. Rather, the query response has been preprocessed and is available for download as a zipped CSV file on GitHub: https://github.com/x-atlas-consortia/hra-pop/blob/main/output-data/v0.11.1/reports/hra/asctb-records.csv.zip.

An exemplary Jupyter Notebook on how to run a SPARQL query against the HRA KG is at https://cns-iu.github.io/hra-kg-supporting-information/#notebook-to-query-the-hra-knowledge-graph-kg.

### Retrieve Mean B Expression Values for CT

The hra-pop *graph* DO (https://lod.humanatlas.io/graph/hra-pop/latest) represents the HRApop dataset and contains mean B expression values for CT inside high-quality experimental datasets. To compute mean B expressions, scanpy^69^, numpy^70^, and anndata^71^ are used. Concretely, scanpy’s to *rank_gene_groups()* method assigns mean B expressions. We normalize gene names with a lookup table from Ensembl Release 111^34^ (https://www.ensembl.org/index.html) to HGNC v2023-09-18^72^ (https://www.genenames.org). Both the code to compute mean B expressions and the look-up table from Ensembl to HGNC are linked in **Supplemental Table 1** under the entry for the hra-pop *graph*.

**Retrieve mean B expression values for a given CT across organs:** Retrieve all experimental datasets from HRApop that contain a canonical CT plus B expression values with this query:

- https://apps.humanatlas.io/api/grlc/hra-pop.html#get-/datasets-with-ct

**Returns**: A table with atlas datasets that contain the given CT. There is one CT-B expression per row. The response includes the dataset source (which portal it was downloaded from), the dataset ID, which must be an Internationalized Resource Identifier (IRI, https://www.w3.org/International/O-URL-and-ident.html), the organ, the donor sex, the tool that assigned the CT, the CT ontology ID (same for every row as provided by user), a human-readable CT label, the number of cells of the given type in the dataset, a B ontology ID, and finally, the mean expression value for all B that characterize this CT computed via CTann tools in HRApop^1^ (see **Box 1**). If multiple CTann tools were used for the dataset, multiple rows are provided to show the different CT and mean B.

### Predict Cell Type Populations and Locations

Seven HRA user stories^1^ have been identified based on interviews with more than 40 atlas architects working on human atlases. While all user stories are directly supported by HRA UIs and the HRA API for users with little to no programming experience, the HRA KG can also be queried directly in support of these user stories. **Supplemental Table 7** lists sample queries that help experts retrieve knowledge from DOs, identify processed data of interest, and run analyses to answer biomedical questions. Here, we detail the queries that support the US#1 (Predict cell type populations) and #2 (Predict spatial origin of tissue samples).

#### Predict cell type populations (US#1)

The HRA KG is used access the hra-pop *graph* (https://lod.humanatlas.io/graph/hra-pop/latest) with cell type populations for AS, datasets, and 3D extraction sites to improve the accuracy of annotations for sc-transcriptomics and sc-proteomics datasets. A demonstration application is available at https://apps.humanatlas.io/us1.

The user provides an extraction site using the RUI. The extraction site is then posted to the HRA API (https://apps.humanatlas.io/api/#post-/hra-pop/rui-location-cell-summary), which returns a predicted cell summary given the 3D collisions of the extraction site with AS inside the 3D Reference Object for which the extraction site was defined. In the background, the API queries a graph inside the HRA KG (https://cdn.humanatlas.io/digital-objects/graph/hra-pop/v0.11.1/assets/atlas-as-cell-summaries.jsonld) to retrieve a predicted cell summary for the extraction site using the SPARQL query at https://github.com/x-atlas-consortia/hra-api/blob/main/src/library/hra-pop/queries/as-weighted-cell-summaries.rq.

#### Predict spatial origin of tissue samples (US#2)

For the inverse case, the cell type populations from the hra-pop *graph* mentioned above can be used to predict the 3D location of datasets with unknown spatial origin. A demonstration application is available at https://apps.humanatlas.io/us2.

The user provides a cell summary for a given dataset whose origin is uncertain or unknown (beyond basic metadata such as the organ and the CTann tool used to assign the CT in the cell summary). The application features an optional dropdown menu to select the organ and tool. Supported organs and tools for the dropdown menu are available via the HRA API endpoints at https://apps.humanatlas.io/api/hra-pop/supported-organs and https://apps.humanatlas.io/api/#get-/hra-pop/supported-tools, respectively. These endpoints run SPARQL queries under the hood: https://github.com/x-atlas-consortia/hra-api/blob/main/src/library/hra-pop/queries/supported-reference-organs.rq (to get the reference organs supported by HRApop) and https://github.com/x-atlas-consortia/hra-api/blob/main/src/library/hra-pop/queries/supported-tools.rq (to get the tools).

Once the user provides the cell type population of the dataset of unknown spatial origin, the application posts this population to an endpoint in the HRA API (https://apps.humanatlas.io/api/hra-pop/cell-summary-report), which takes a minimum of a CT ID and a column for the percentage of that CT in the dataset. In the background, the HRA API runs the SPARQL query at https://github.com/x-atlas-consortia/hra-api/blob/main/src/library/hra-pop/queries/select-cell-summaries.rq, which returns a listing of the most similar AS, datasets, and extraction sites (by cosine similarity).

## Code Availability

All the code used to construct and deploy the HRA KG v2.2 is available on GitHub at https://github.com/hubmapconsortium/hra-kg and URLs are provided in **Supplemental Table 4**. Documentation, including an additional overview of HRA KG construction code is provided in the supporting information repository for this paper at https://github.com/cns-iu/hra-kg-supporting-information/?tab=readme-ov-file#github-repositories-used-for-hra-kg-construction. We also created documentation with annotated screenshots to show how to run pre-made SPARQL queries against the HRA KG via https://grlc.io, see https://cns-iu.github.io/hra-kg-supporting-information/#how-to-run-queries-via-our-openapi-spec.

All code was released under the MIT License.

## Author Contributions

A.B. led the writing of the paper, built the companion website at https://cns-iu.github.io/hra-kg-supporting-information, and leads the HRApop effort. With B.W.H. and K.B, he shares corresponding authorship. B.W.H and J.H. built the hra-do-processor and engineered the CCF Ontology and HRA data structures. B.W.H. leads the development of the HRA user interfaces and the generation and publication of the HRA KG. A.B. and B.W.H. compiled a notebook to show basic usage of the HRA KG on the companion website. E.M.Q. compiled CTann annotation crosswalks used in and published with the HRA KG. M.M. oversees HRA ontology engineering work. K.B. leads the HRA effort and specified HRA KG usage to help focus HRA KG development and documentation. A.B., B.W.H., J.H., E.M.Q., and K.B. wrote the paper. All other authors reviewed and commented on the paper.

## Acknowledgments

The authors would like to thank Sören Auer, Ino de Bruijn, Yongqun (Oliver) He, Nancy Ruschman, James McLaughlin, Deanne Taylor, and Maria-Esther Vidal for their expert comments and suggestions on an earlier version of this paper. Libby Maier supported the design of figures.

The HRA is under active development by HuBMAP, SenNet, KPMP, GUDMAP, and the National Institute of Diabetes and Digestive and Kidney Diseases (NIDDK) with expert input by the HRA Editorial Board and in close collaboration with experts from more than 15 other consortia. K.B. is a co-director of and is funded by the CIFAR MacMillan Multiscale Human program.

This research has been supported by the NIH Common Fund through the Office of Strategic Coordination/Office of the NIH Director under awards:

- OT2OD033756 and OT2OD026671 (A.B., B.W.H., J.H., E.M.Q., M.M., K.B.);
- OT2OD026675 and OT2OD033759 (A.B., NIH JumpStart Award/Fellowship)
- OT2OD030545 (A.B., B.W.H., K.B.)

Further, this work was supported by:

- the SenNet Consortium Organization and Data Coordinating Center (CODCC) under award number U24CA268108-01 (A.B., B.W.H., E.M.Q., K.B.);
- by the NIDDK under award U24DK135157 (B.W.H., K.B.);
- by the KPMP grant U2CDK114886 (A.B., B.W.H., K.B.);
- and the NIH National Institute of Allergy and Infectious Diseases (NIAID), Department of Health and Human Services under BCBB Support Services Contract HHSN316201300006W/HHSN27200002.

This research was supported in part by the Intramural Research Program of the U.S. National Institutes of Health. The funders had no role in study design, data collection and analysis, decision to publish, or preparation of the manuscript. The content is solely the responsibility of the authors and does not necessarily represent the official views of the National Institutes of Health.

## Supplemental Figures

**Supplemental Figure 1.**
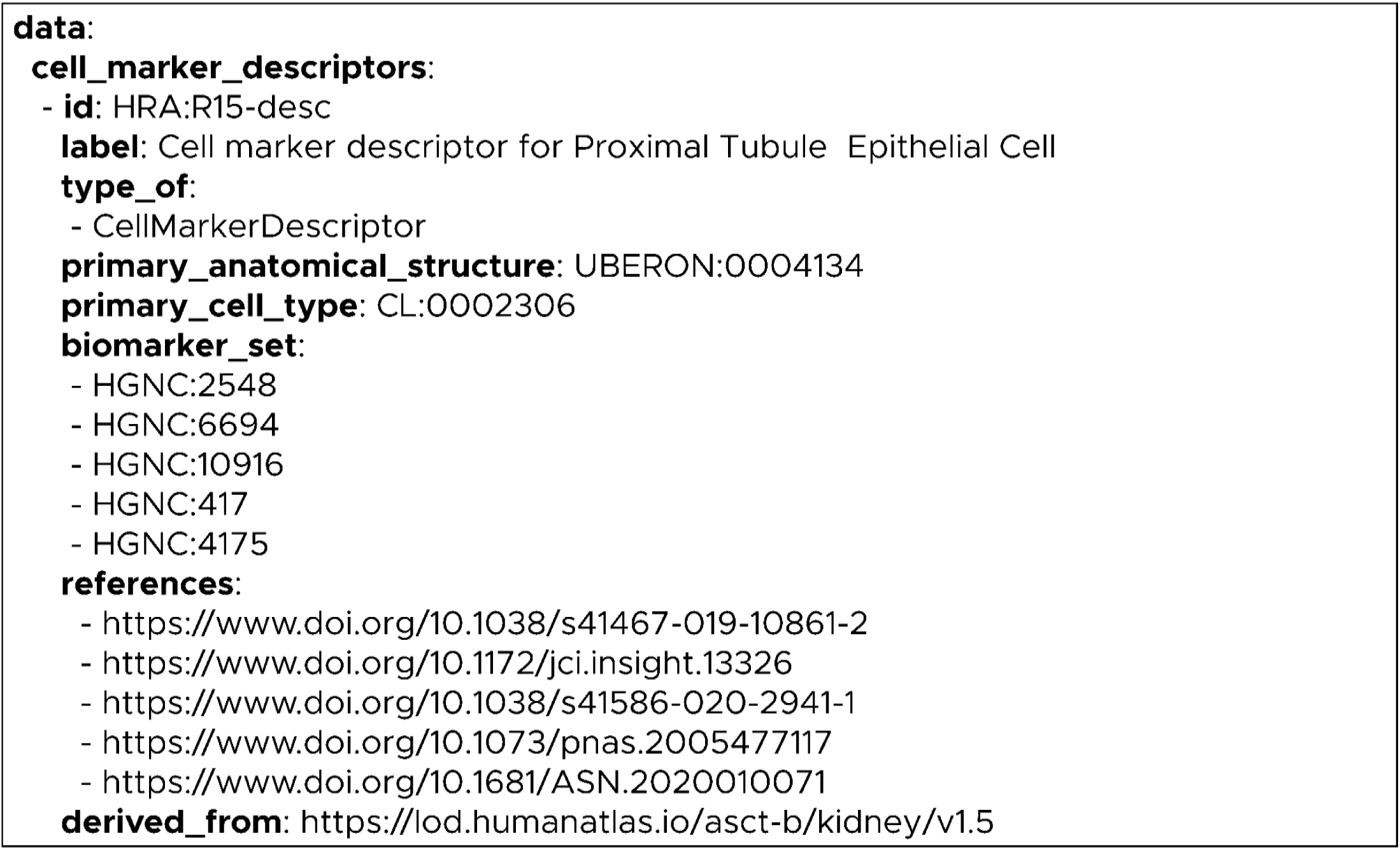
An excerpt of a normalized structure from the ASCT+B table for the kidney, with a focus on *cell_marker_descriptors*, where primary CT, primary anatomical location, and associated characterizing B are detailed, and where references to support the claims regarding the CT and its B are provided. An example in YAML is provided at https://github.com/cns-iu/hra-kg-supporting-information/blob/main/docs/intermediary_format.yaml.

**Supplemental Figure 2.**
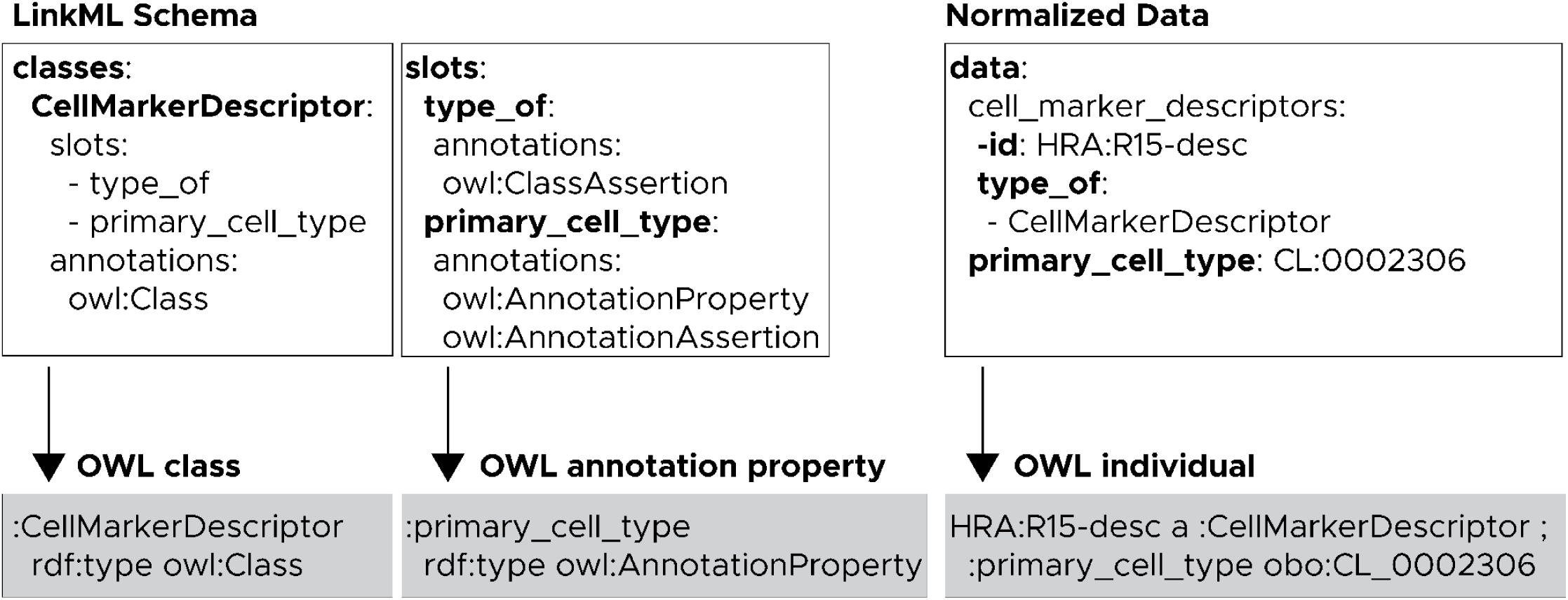
This diagram shows an excerpt of a LinkML schema (represented by the two boxes on the left) and the corresponding acquired data (the box on the right). Arrows illustrate the transformation of the input text into OWL constructs at the bottom. In the schema boxes, classes and slots are directly mapped to OWL classes and OWL properties as specified in the *annotations* field. In the data box, each data item pair is translated into an OWL assertion statement. For example, the data pair *type_of: CellMarkerDescriptor* generates a class assertion that indicates the data object belongs to the CellMarkerDescriptor class. Similarly, the data pair *primary_cell_type: CL:0002306* produces an annotation assertion that tells the same data object identifies the epithelial cell in the proximal tubule of the kidney (CL:0002306) as the primary CT.

## Supplemental Tables

**Supplemental Table 1.**
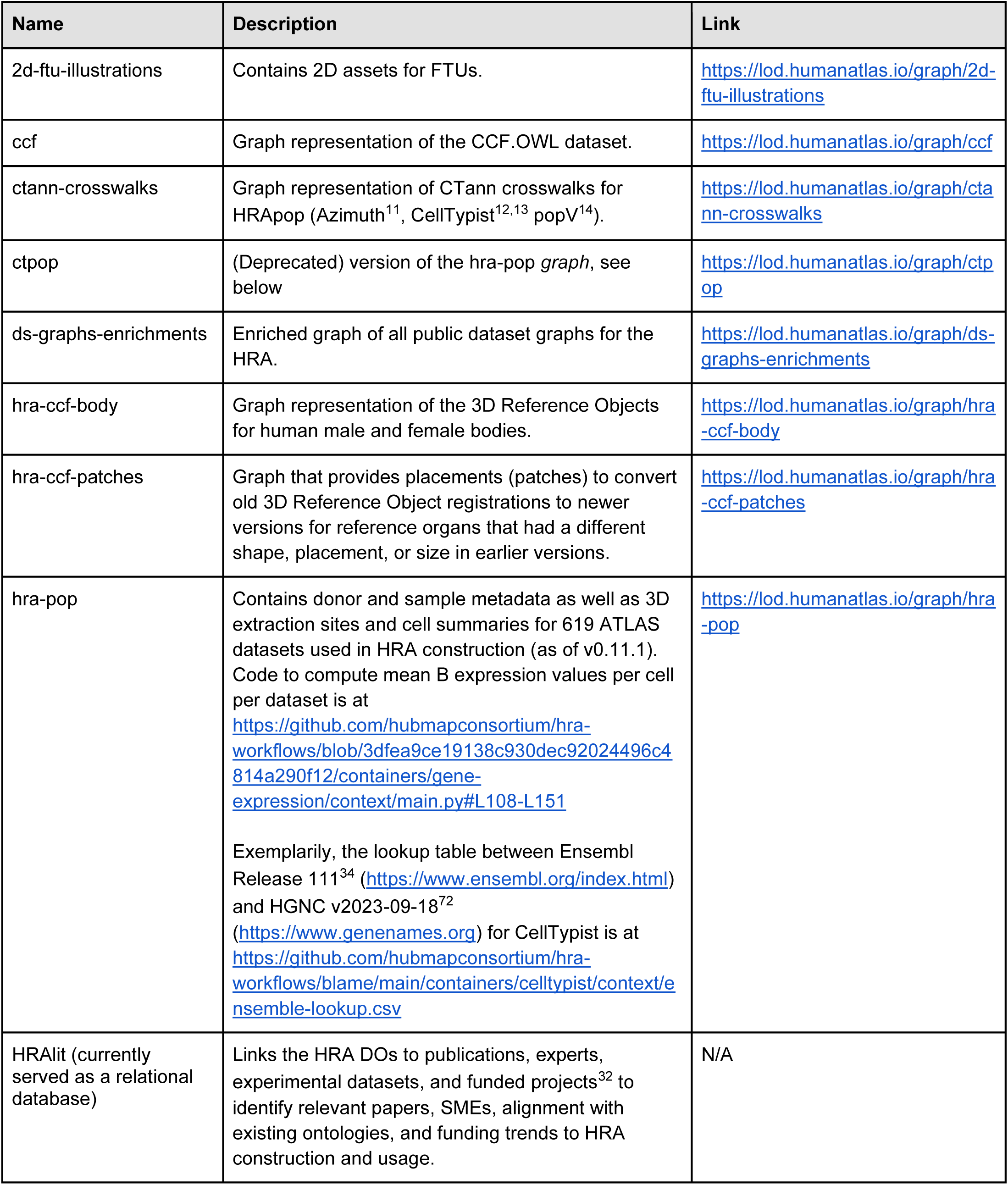
Graphs of the HRA KG (https://lod.humanatlas.io/graph).

**Supplemental Table 2.** Collections of the HRA KG (https://lod.humanatlas.io/collection). All structures are built using standardized terms for labels, which are stored in the specialized CCF vocabulary, see CCF Ontology in Box 1.

| Name | Description | Link |
| --- | --- | --- |
| ds-graphs | Consolidates experiment-result datasets (*ds-graph*) into a single graph data structure. Contains processed graphs of datasets including donor and sample metadata from different consortia and tissue providers registered to HRA via 3D extraction sites, available at https://lod.humanatlas.io/ds-graph. Also present are HRA *graph* DOs (https://lod.humanatlas.io/graph), which are externally processed RDF graphs, such as crosswalks between CT assigned by CTann tools and CT in ASCT+B tables. | https://lod.humanatlas.io/collection/graphs |
| hra | Includes HRA DO types with a DOI. As of HRA v2.2, those are *2d-ftu*, *asct-b*, *ctann*, *omap*, *ref-organ*, and *vascular-geometry* DO types. Other DO types can be added in the future once they have DOIs. | https://lod.humanatlas.io/collection/hra |
| hra-api | Centers on *asct-b* and associated 3D models (*ref-organ*, *landmark*) used in various HRA applications for visualizing and organizing reference data. | https://lod.humanatlas.io/collection/hra-api |
| hra-ols | HRA collection subset that focuses solely on AS, CT, and B (*asct-b* only). It is used by OLS (https://www.ebi.ac.uk/ols4) for validation. The source for the files on the LOD server are available at https://github.com/hubmapconsortium/3d-hra-ref-object-validation. | https://lod.humanatlas.io/collection/hra-ols |

**Supplemental Table 3.**
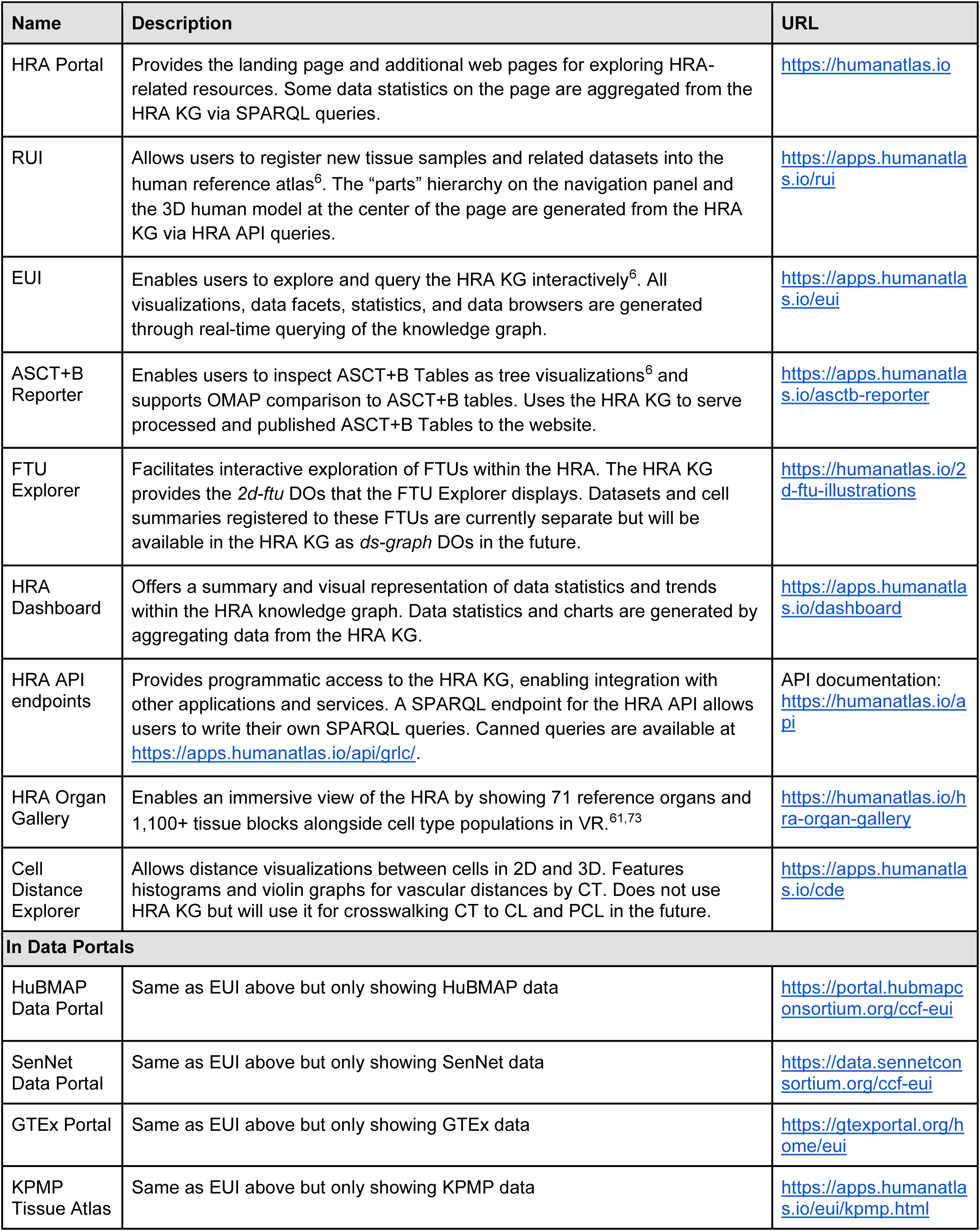
Most HRA applications use the HRA KG as their backend database. Also listed are external applications that use the HRA KG.

**Supplemental Table 4.**
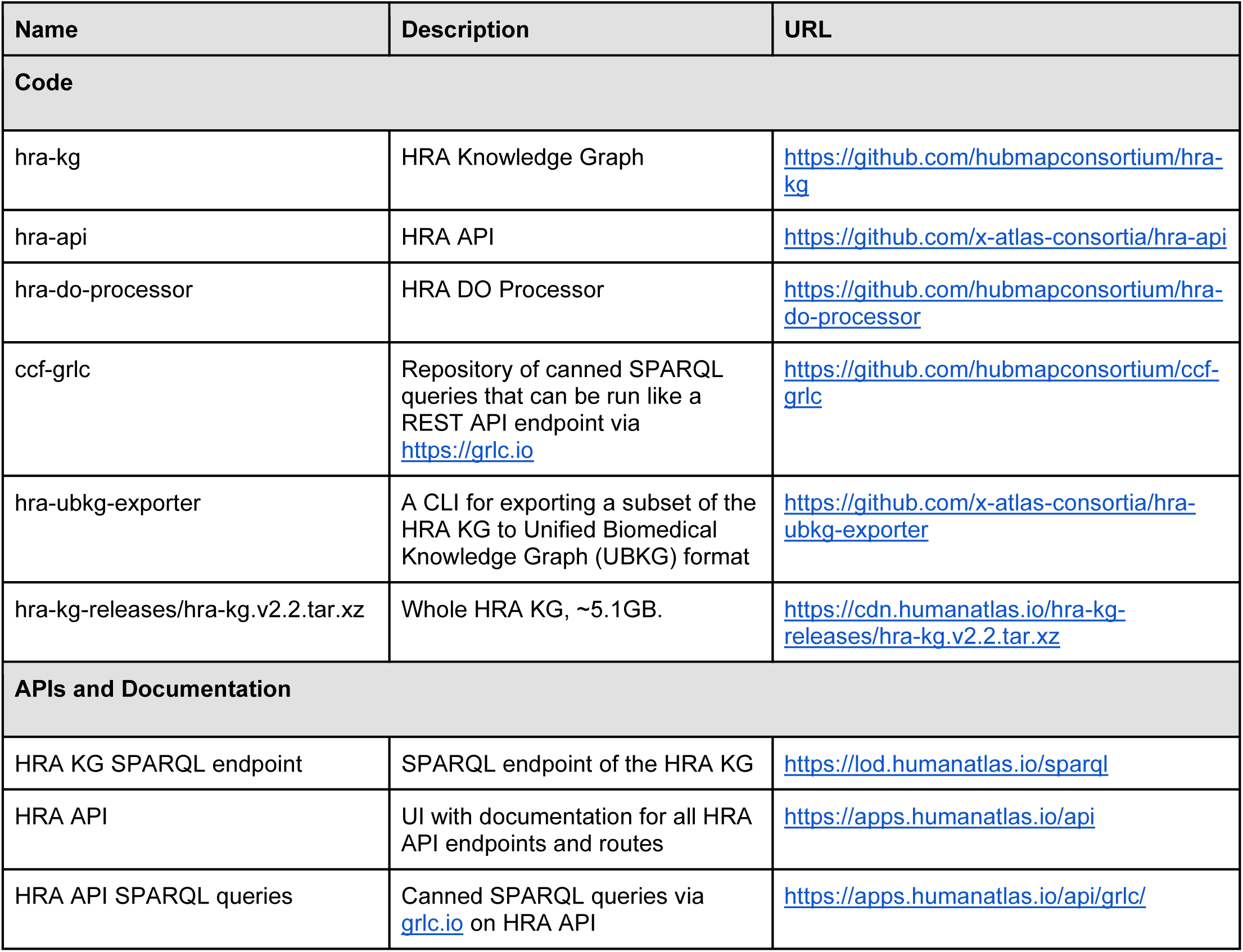
An overview of all GitHub repositories used to construct, deploy, and use the HRA KG.

| Name | Description | URL |
| --- | --- | --- |
| **Code** |  |  |
| hra-kg | HRA Knowledge Graph | https://github.com/hubmapconsortium/hra-kg |
| hra-api | HRA API | https://github.com/x-atlas-consortia/hra-api |
| hra-do-processor | HRA DO Processor | https://github.com/hubmapconsortium/hra-do-processor |
| ccf-grlc | Repository of canned SPARQL queries that can be run like a REST API endpoint via https://grlc.io | https://github.com/hubmapconsortium/ccf-grlc |
| hra-ubkg-exporter | A CLI for exporting a subset of the HRA KG to Unified Biomedical Knowledge Graph (UBKG) format | https://github.com/x-atlas-consortia/hra-ubkg-exporter |
| hra-kg-releases/hra-kg.v2.2.tar.xz | Whole HRA KG, ~5.1GB. | https://cdn.humanatlas.io/hra-kg-releases/hra-kg.v2.2.tar.xz |
| **APIs and Documentation** |  |  |
| HRA KG SPARQL endpoint | SPARQL endpoint of the HRA KG | https://lod.humanatlas.io/sparql |
| HRA API | UI with documentation for all HRA API endpoints and routes | https://apps.humanatlas.io/api |
| HRA API SPARQL queries | Canned SPARQL queries via grlc.io on HRA API | https://apps.humanatlas.io/api/grlc/ |

**Supplemental Table 5.** List of LinkML schemas in the HRA KG.

| Schema Description | Link |
| --- | --- |
| Graph schema for ASCT+B tables | https://github.com/hubmapconsortium/hra-do-processor/blob/main/schemas/src/digital-objects/asct-b.yaml |
| Graph schema for 3D Reference Objects for organs | https://github.com/hubmapconsortium/hra-do-processor/blob/main/schemas/src/digital-objects/ref-organ.yaml |
| Graph schema for 2D FTUs | https://github.com/hubmapconsortium/hra-do-processor/blob/main/schemas/src/digital-objects/2d-ftu.yaml |
| Graph schema for the 3D Landmarks | https://github.com/hubmapconsortium/hra-do-processor/blob/main/schemas/src/digital-objects/landmark.yaml |
| Graph schema for OMAPs | https://github.com/hubmapconsortium/hra-do-processor/blob/main/schemas/src/digital-objects/omap.yaml |
| Graph schema for experimental dataset metadata | https://github.com/hubmapconsortium/hra-do-processor/blob/main/schemas/src/digital-objects/ds-graph.yaml |
| Graph schema for external graph files (metadata only) | https://github.com/hubmapconsortium/hra-do-processor/blob/main/schemas/src/digital-objects/graph.yaml |
| Graph schema for vocabulary resources (metadata only) | https://github.com/hubmapconsortium/hra-do-processor/blob/main/schemas/src/digital-objects/vocab.yaml |
| Graph schema for curated graph collections (metadata only) | https://github.com/hubmapconsortium/hra-do-processor/blob/main/schemas/src/digital-objects/collection.yaml |
| Graph schema for cell summaries | https://github.com/hubmapconsortium/hra-do-processor/blob/main/schemas/src/modules/cell-summary.yaml |
| Graph schema for spatial data | https://github.com/hubmapconsortium/hra-do-processor/blob/main/schemas/src/modules/spatial.yaml |
| Graph schema for spatial collision data | https://github.com/hubmapconsortium/hra-do-processor/blob/main/schemas/src/modules/collision.yaml |
| Graph schema for spatial corridor data | https://github.com/hubmapconsortium/hra-do-processor/blob/main/schemas/src/modules/corridor.yaml |
| Graph schema for assay dataset metadata | https://github.com/hubmapconsortium/hra-do-processor/blob/main/schemas/src/modules/dataset.yaml |
| Graph schema for donor metadata. | https://github.com/hubmapconsortium/hra-do-processor/blob/main/schemas/src/modules/donor.yaml |
| Graph schema for sample metadata | https://github.com/hubmapconsortium/hra-do-processor/blob/main/schemas/src/modules/sample.yaml |
| Graph schema for defining entities | https://github.com/hubmapconsortium/hra-do-processor/blob/main/schemas/src/shared/entity-base.yaml |
| Graph schema for defining instances | https://github.com/hubmapconsortium/hra-do-processor/blob/main/schemas/src/shared/instance-base.yaml |
| Graph schema for defining graph metadata | https://github.com/hubmapconsortium/hra-do-processor/blob/main/schemas/src/shared/metadata-base.yaml |

**Supplemental Table 6.**
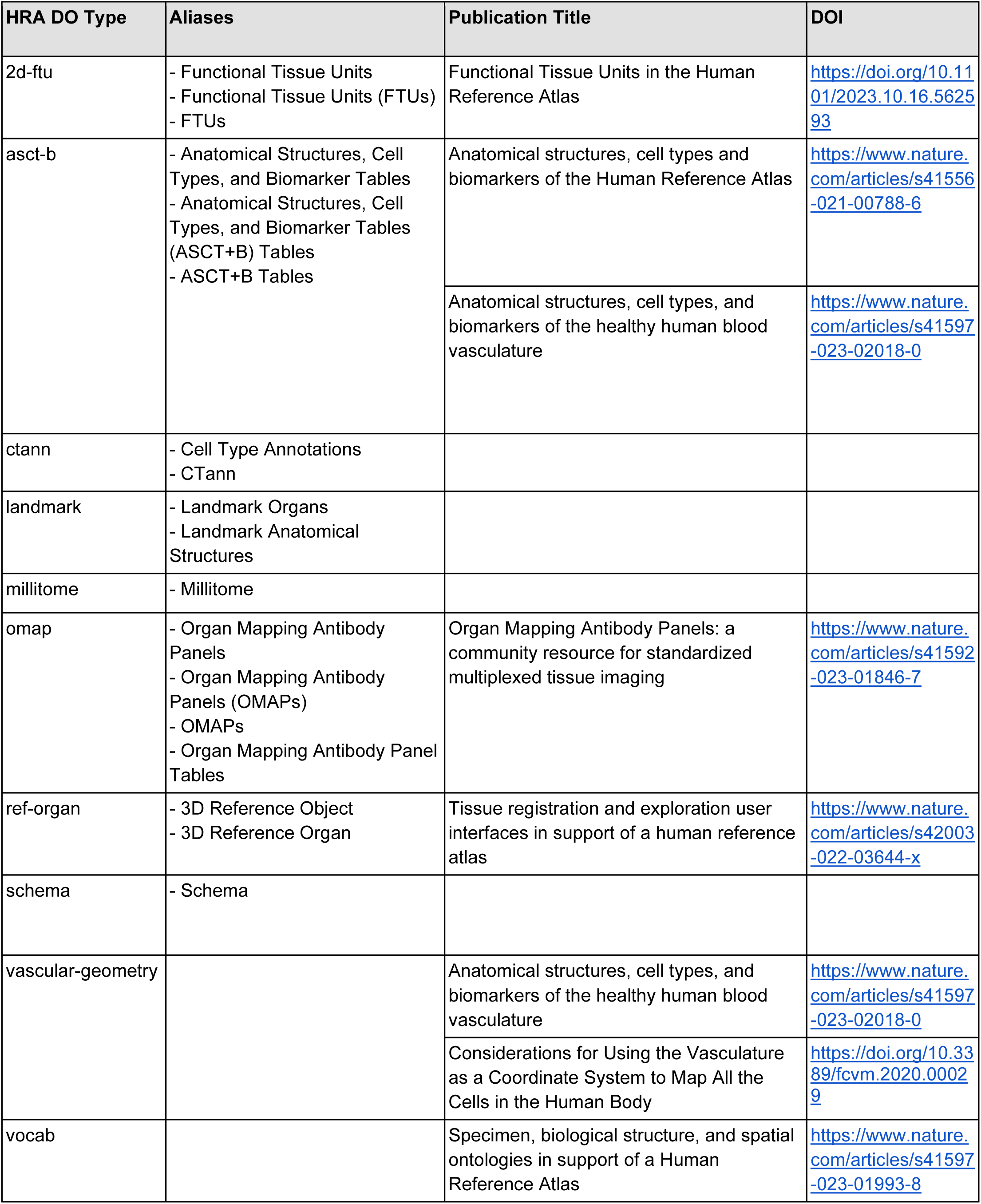
Publications on HRA DOs plus aliases used throughout HRA publications and applications.

**Supplemental Table 7.**
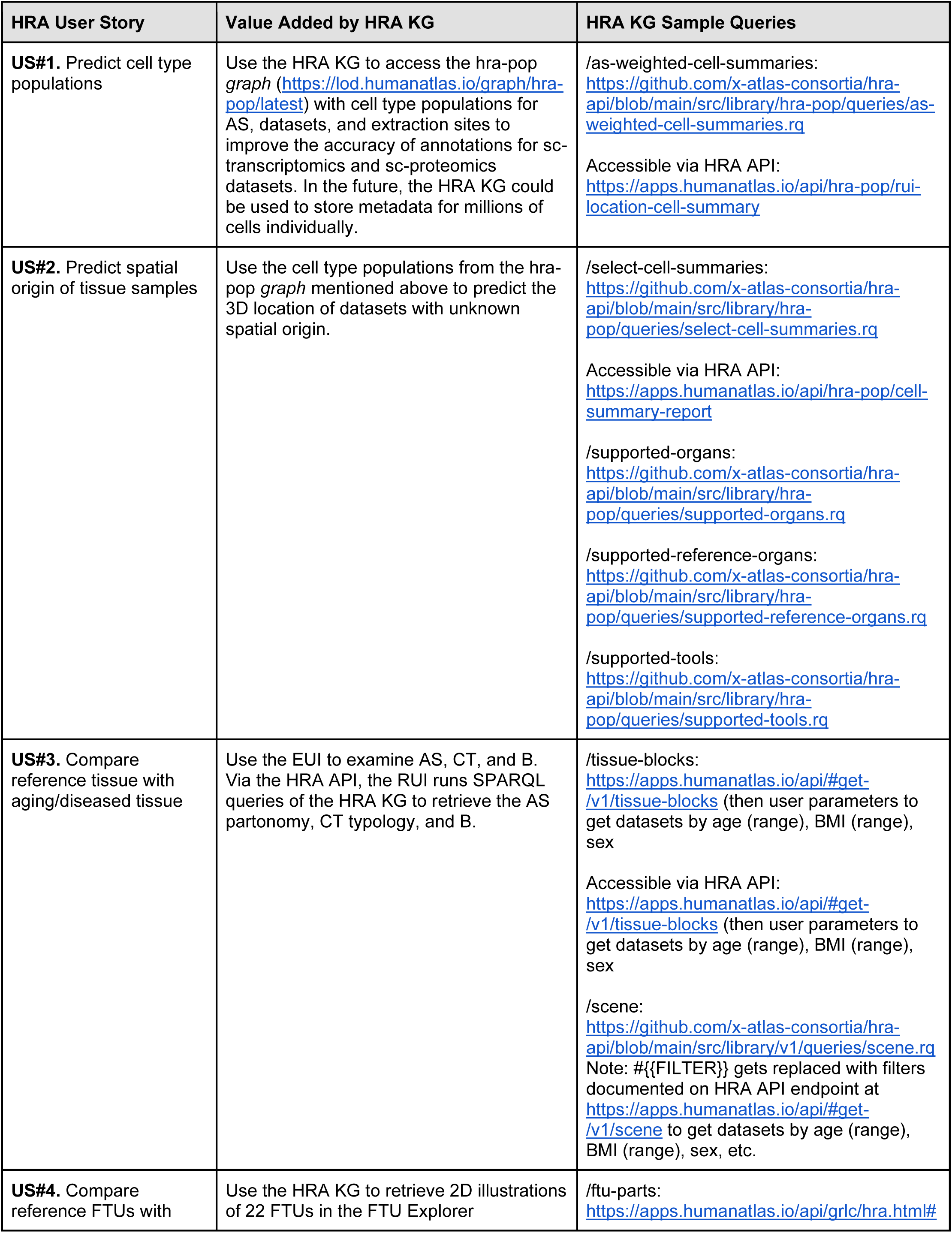
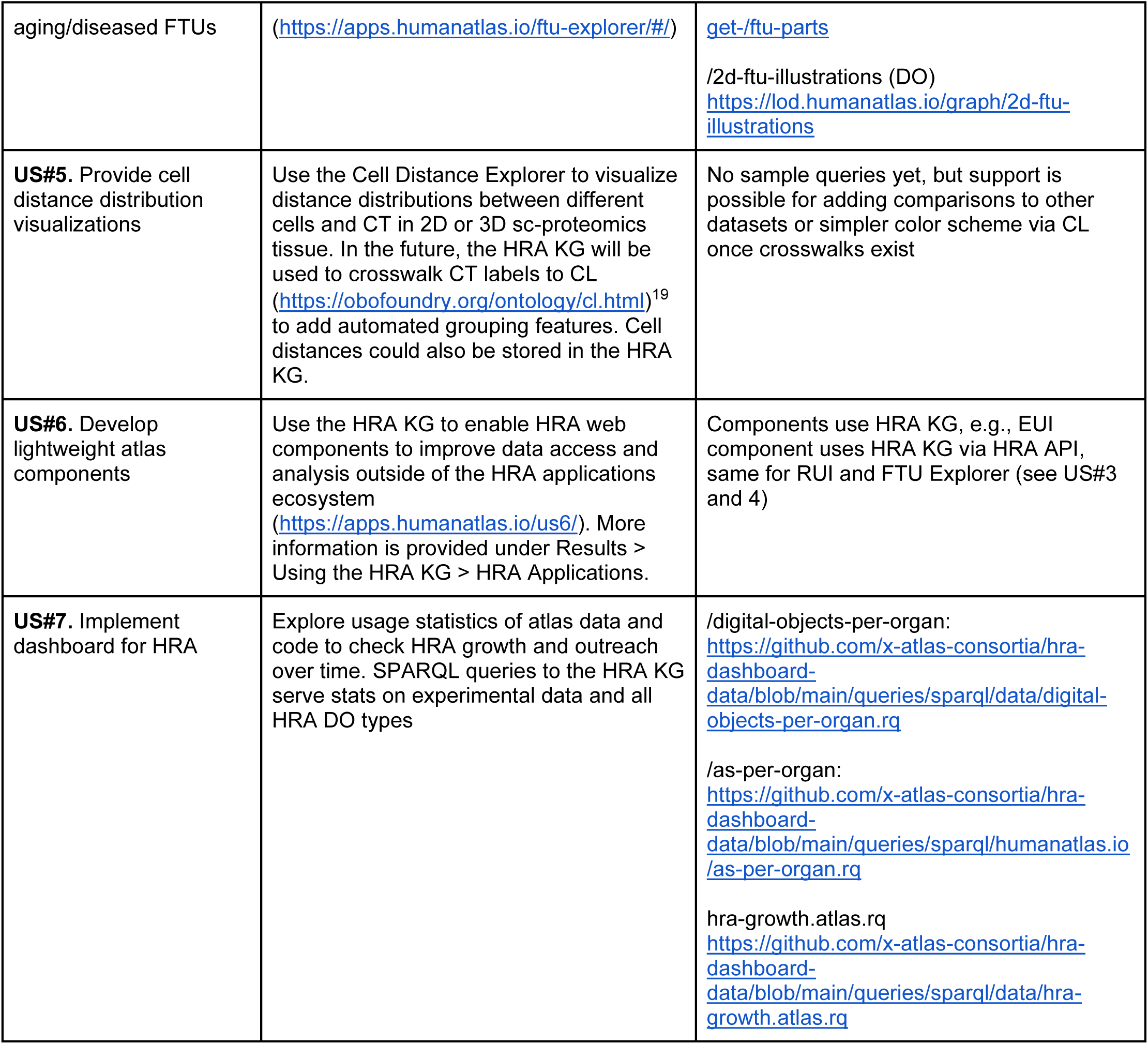
HRA User Stories and how the HRA KG supports them.

